# EZH1/2 inhibition improves immunotherapy response through MHC Class II de-repression and neutrophil reprogramming

**DOI:** 10.64898/2026.06.01.725956

**Authors:** Avery R. Childress, Dave-Preston Esoe, Xiulong Song, Christain M. Gosser, Yindan Lin, Daniel R. Plaugher, Tanner J. DuCote, Kassandra J. Naughton, Erika M. Skaggs, Harrison Yang, Ryan Goettl, Jinpeng Liu, Zhonglin Hao, Albert E. Fliss, Daisuke Honma, Todd Burus, Feitong Lei, Bin Huang, Ellen J. Beswick, Christine F. Brainson

## Abstract

Squamous cell carcinoma of the lung is a difficult-to-treat cancer with high prevalence in the US, and particularly in Kentucky. Here, we sought to test if the EZH1/2 inhibitor valemetostat improves anti-PD1 responses in squamous cell lung cancer models, and to develop ex vivo models to test immunotherapy drug combinations. We found that valemetostat augmented anti-tumor responses to anti-PD1 therapy, led to up-regulation of tumor cell specific Major Histocompatibility Complex Class II (MHC Class II), and drove systemic neutrophil maturation. Likewise, *Ezh2* knock-out mice produced neutrophils that were more apoptotic, less migratory, and less able to produce extracellular nets, but had similar ability to kill bacteria as *Ezh2*-WT neutrophils. To test tumor responses to differing neutrophil populations, we engineered three-dimensional air-liquid interface cultures with tumoroids, lung mesenchymal cells, and T cells, with and without bone marrow containing neutrophils and myeloid progenitors from distinct donors. Bone marrow from tumor-naïve or mice with actively growing untreated tumors boosted tumoroid growth, while bone marrow from spontaneously tumor-rejected or mice with tumors treated with valemetostat was anti-tumor. MHC Class II blockade lowered the ability of bone marrow to boost tumor growth, and reduced the ability of valemetostat with anti-PD1 to reduce tumoroid growth. Patient samples revealed a strong negative correlation between EZH2 and MHC Class II, suggesting that targeting EZH2 activity could lead to marked increase in MHC Class II and improve treatment responses in lung squamous cell carcinomas.

## INTRODUCTION

Immunotherapies, including those that target the PD1/PD-L1 immune checkpoint, have revolutionized lung cancer treatment. However, only 18% of lung squamous cell carcinoma patients have a durable response to immunotherapies^1–3^. Lung squamous cell carcinomas fall under the larger umbrella of non-small cell lung cancer (NSCLCs), and are characterized by malignant cells organized in a fully stratified squamous epithelium, and high proportions of tumor-associated neutrophils^4–6^. Given the common use of immunotherapy to treat lung squamous cell carcinomas, new treatments are needed to improve immunotherapy efficacy in this patient population.

The Polycomb Repressive Complex 2 contains the histone methyltransferase EZH2, or the closely related protein EZH1, that can add the second and third methyl groups to the lysine at the 27^th^ amino acid position of the histone H3 tail^7^. This epigenetic modification, termed H3K27me3, leads to transcriptional silencing of numerous genes, including transcripts encoding antigen presentation and immune activation proteins^8–11^. Several ATP-competitive EZH2 inhibitors have been developed to date. GSK126, while unsuccessful in clinical trials, remains a valuable tool compound, and tazemetostat (EPZ6438) was FDA-approved for epithelioid sarcoma and follicular lymphoma administered but was recently removed from the market^12,13^. More recently, the dual EZH1/2 inhibitor valemetostat was developed. Valemetostat is administered orally once daily at 200 mg, one eighth of the total daily dose of tazemetostat, and is currently approved in Japan for use in relapsed/refractory adult T cell leukemia and peripheral T cell lymphoma^14,15^.

Numerous studies have tested the combined effects of EZH2 inhibition with anti-PD1 therapy, and several clinical trials have emerged from the promising pre-clinical data. Major findings include an increase in activated CD8 T cells and an increase in MHC Class II expression on tumor cells^10,11^. Although MHC Class II expression is typically associated with professional antigen presenting cells, including macrophages and dendritic cells, numerous epithelial cell types in the lung have the capacity to express MHC Class II^16,17^. In particular, lung squamous cell carcinomas originate from lung epithelial cell lineages, including club cells and basal cells, capable of inducible expression of MHC class II^16,17^, suggesting that reactivation of this pathway may enhance antigen presentation and improve responsiveness to ICI.

Recapitulating the microenvironment *ex vivo* provides a reproducible and tunable model to study the complex cellular interactions that occur within the TME. Specifically, studying ICIs *in vitro* remains challenging given that current systems typically require fresh tumor tissue from a murine or human host^18–20^. Some studies have focused on combining only tumor cells with T cells, particularly with Chimeric-Antigen Receptor (CAR) T cells with cells engineered to, or naturally expressing, the CAR antigen^21,22^. While these reductionist systems offer valuable mechanistic insight, they do not fully capture the dynamic tumor–immune interactions present in an intact tumor ecosystem. Using our murine models, we allow established tumoroids to shape the immune cell landscape, which can then be modulated through the addition of our therapeutic interventions. These cultures can be further augmented by varying immune cell ratios, incorporating immune cells from distinct backgrounds (tumor-bearing versus non-tumor-bearing), and blocking specific signaling pathways to assess how these factors influence therapeutic response.

## RESULTS

Kentucky is a hot spot to study lung cancer, having the highest rates of lung and bronchus cancers relative to the rest of the United States (**Figure 1A**). Furthermore, using histology codes for the most common types of non-small cell lung cancer: adenocarcinoma (ADC), squamous cell carcinoma (SCC), and mixed/undifferentiated lung cancers (Mixed/Not Otherwise Specified aka NOS), Appalachia Kentucky has the highest percentages of SCCs (34%) and mixed/undifferentiated cancers (28%), which also have poor prognosis (**Figure 1B**). Together, these data indicate an immediate need to understand how to improve outcomes for these patients. Given that anti-PD1/PD-L1 targeting immunotherapies are the most commonly prescribed treatments for advanced stage NSCLCs, we set out to test if adding an epigenetic EZH1/2 inhibitor, valemetostat, can improve treatment outcomes in preclinical models. To this end, we utilized a syngeneic graft model that recapitulates the histology and immune cell infiltration observed in primary tumors ^11^. Tumor bearing mice were randomized into four treatment arms: placebo, valemetostat tosylate (135 mg/kg, oral gavage QD), anti-PD1 antibody (7mg/kg, intraperitoneal injection 3x/week), or the combination treatment (**Figure 1C**). Drugs were well tolerated with slight weight loss observed in all groups likely due to daily oral gavage (**Supp. Figure 1A**). While both single-agent treatments slowed tumor growth, tumor regression was observed only in the combination group, with significant tumor reductions observed from day 20 of treatment (**Figure 1D&E, Supp. Figure 1B**).

**Figure 1:**
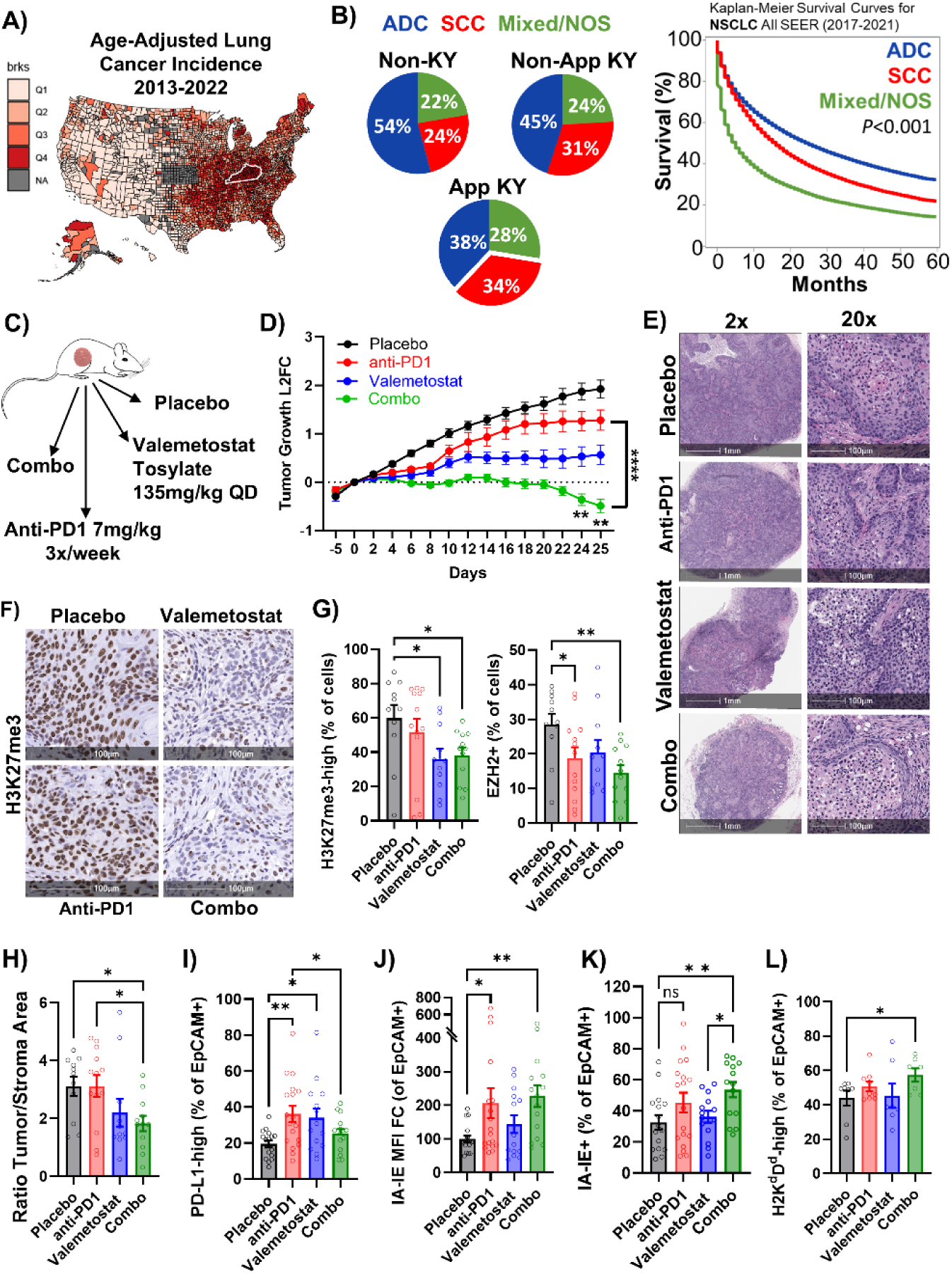
Valemetostat increases anti-PD1 efficacy and tumor MHC Class II expression. A) Heat map of age-adjusted lung cancer incidence complied by county from 2013-2022. **B)** Data compiled from the SEER database to show proportions of lung cancer histological subtypes within Kentucky and in the other SEER databases, and overall survival of each cohort. **C)** Schematic of mouse cohorts and treatments. **D)** Tumor growth from the syngeneic mouse model with indicated treatments. ** indicates p<0.003 by two-way ANOVA for every pair-wise comparison with multiple comparisons and Holm-Šídák’s *post-hoc* test from day 22 till end, anti-PD1 vs combo comparison is ***p<0.0001 from day 11 onward. **E)** Tumors from mice were removed and sectioned then stained using Hematoxylin. Images at 2x and 20x magnification are shown. **F)** Immunohistochemistry for H3K27me3, scale bars = 100µm. **G)** H3K27me3+ cells and EZH2+ cells quantified by HALO on IHC stained sections, for all graphs mean +/- SEM is plotted. **H)** Ratio of tumor area to stroma area by machine learning algorithm. **I-L)** EpCAM+ tumor cells were gated for **(I)** % PD-L1-high **(J)** % IA-IE+ **(K)** Fold change in IA-IE MFI **(L) %** H2K^d^D^d^-high, *<0.05, **<0.01 by one-way ANOVA with pairwise comparisons. Also see Supp. Figure 1.

To measure drug responses in tumors, after 21 days of treatment the remaining lesions were harvested for histological and flow cytometric analyses. Immunohistochemistry revealed a decrease in both EZH2 and H3K27me3 in combination treated tumors (**Figure 1F&G, Supp. Figure 1C**). Given that EZH2 is an E2F target gene^23^, the reduction in EZH2 is likely a result of decreased proliferation rather than destabilization of EZH2 by valemetostat. Using a computational algorithm to quantify areas of malignant epithelial cells versus area that were acellular or predominately immune infiltrates, defined as stroma, we found that combination-treated tumors demonstrated reduced tumor-to-stroma area ratios (**Figure 1H, Supp. Figure 1D**). Next, we utilized flow cytometry on dissociated tumors to analyze cell markers present on the EpCAM+ tumor cells (**Supp. Figure 1 E&F**). PD-L1 expression was increased in all treatment groups relative to placebo, though levels were significantly lower in the combination treatment group compared to anti-PD1 monotherapy (**Figure 1I**). Given that MHC class I and II expression have been shown to be directly regulated by EZH2^8,11^, we next evaluated these markers and found these markers to be highest in the combination treatment group (**Figure 1J-L, Supp. Figure 1G**).

We next sought to determine how valemetostat treatment alters the cellular composition of the tumor microenvironment (TME). First, using H&E-stained sections from harvested tumors, we performed nuclear profiling with an artificial intelligence trained algorithm^24^. As previously observed^11,25^, neutrophils were the largest immune infiltrate in this model (**Figure 2A**), and there were reductions in macrophage and tumor cell proportions in combination treated tumors (**Figure 2B**). Given these shifts in the myeloid and tumor compartments, we next evaluated whether combination therapy also modulated T cell activation within the TME. We found that the percentages of IFNγ-expressing CD3+ T cells were significantly increased in the combination treatment group (**Figure 2C, Supp. Figure 2A**). Additionally, PD1-high T cells were increased, further suggesting an increased T cell activation in combination treated tumors (**Figure 2D**). We next examined CD8+ T cells and observed increased proportions of both IFNγ+ and PD1-high cells in this population as well (**Figure 2E&F**). For canonical neutrophil markers, CD11b and Ly6G, there was a trending decrease in the combination group that did not reach statistical significance, and neutrophils remained a very high percentage of CD45+ cells regardless of treatment (**Figure 2G, Supp. Figure 2B**). Overall, there was a small but significant decrease in the myeloid marker CD11b+, mirroring an increase in CD11b- negative populations (**Figure 2 H&I**). Collectively, these findings suggest that combination treatment with anti-PD1 and valemetostat promotes more activated T cells and fewer myeloid cells within the tumor microenvironment. However, it is important to note that neutrophils remained the predominant cell type in the TME of these lung squamous cell carcinoma grafts.

**Figure 2:**
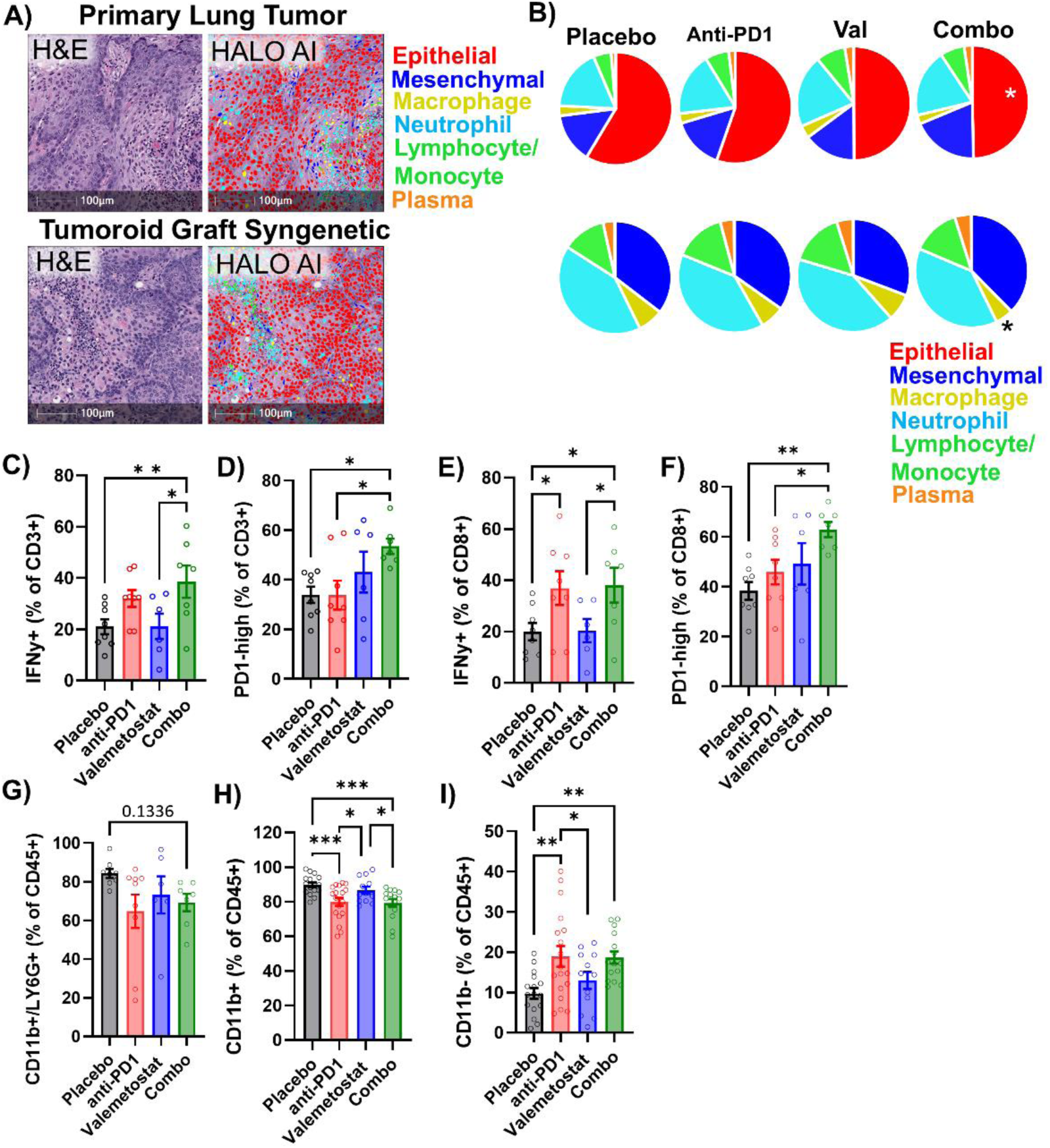
Valemetostat increases lymphoid activation and decreases myeloid proportions. **A)** Images of the original primary autochthonous *Lkb1/Pten* tumor and a syngeneic graft seeded from tumoroids from the original mouse. H&E and HALO nuclear profiler images are show, scale bars = 100µm. Legend indicates cell types identified by nuclear profiler. **B)** Pie charts depicting proportions of different cell types as quantified with the nuclear phenotype. Including epithelial/tumor cells is shown on top, and only non-epithelia/tumor cells is shown on bottom, * p<0.05 by two-tailed t-test. **(C-I)** Flow cytometry from 2 separate mouse cohorts. Indicated populations were gated for: **(C)** % IFNy+ T cells **(D)** % PD1-high T cells **(E)** % IFNy+ CD8+ T cells **(F)** % PD1-high CD8+ T cells **(G)** % CD11b+/LY6G+ **(H+I)** % CD11b+ and CD11b- from two flow stains per tumor. For all graphs mean +/- SEM is plotted, *<0.05, **<0.01, ***<0.001 by one-way ANOVA with pairwise comparison. See also Supp. Figure 2.

The differentiation stage of neutrophils can be measured by assessing their nuclear morphology from a solid bean-shaped nucleus at the myelocyte stage to a round nuclear with a hole in the center in the banded stage to a multi-lobed nuclear in the hyper-segmented stage (**Figure 3A**). From bone marrow, both high-density more mature neutrophils and low-density neutrophils as well as other immune cell types can be enriched using a dual histopaque gradient (**Supp. Figure 3A&B**). In stained cytospins of high-density enriched neutrophils, valemetostat- and combination-treated mice exhibited markedly higher proportions of mature and hyper-segmented neutrophils compared to controls (**Figure 3B**). Using published gene signatures associated with early, intermediate, and late stages of neutrophil differentiation^11,26,27^, we assessed treatment-induced transcriptional changes using RNA-sequencing of low-density neutrophils. Gene set enrichment analysis (GSEA)^28^ revealed that bone marrow from mice treated with valemetostat alone or in combination showed enrichment of genes linked to late-stage neutrophil differentiation when compared to placebo-treated bone marrow (**Figure 3C**). Furthermore, established gene signatures revealed that valemetostat and combo treated bone marrow had increased gene expression of genes involved in chromatin organization and genes repressed by H3K27me3 (**Figure 3D**, **Supp. Table 1**). Additionally, gene signatures related to myeloid differentiation, lipid metabolism, and senescence were upregulated following treatment. Antimicrobial and humoral defense pathways were also enhanced, suggesting that treated neutrophils retain their capacity for bacterial defense. Conversely, MYC target genes, as well as pathways associated with DNA damage, proliferation, and migration, were downregulated with valemetostat.

**Figure 3:**
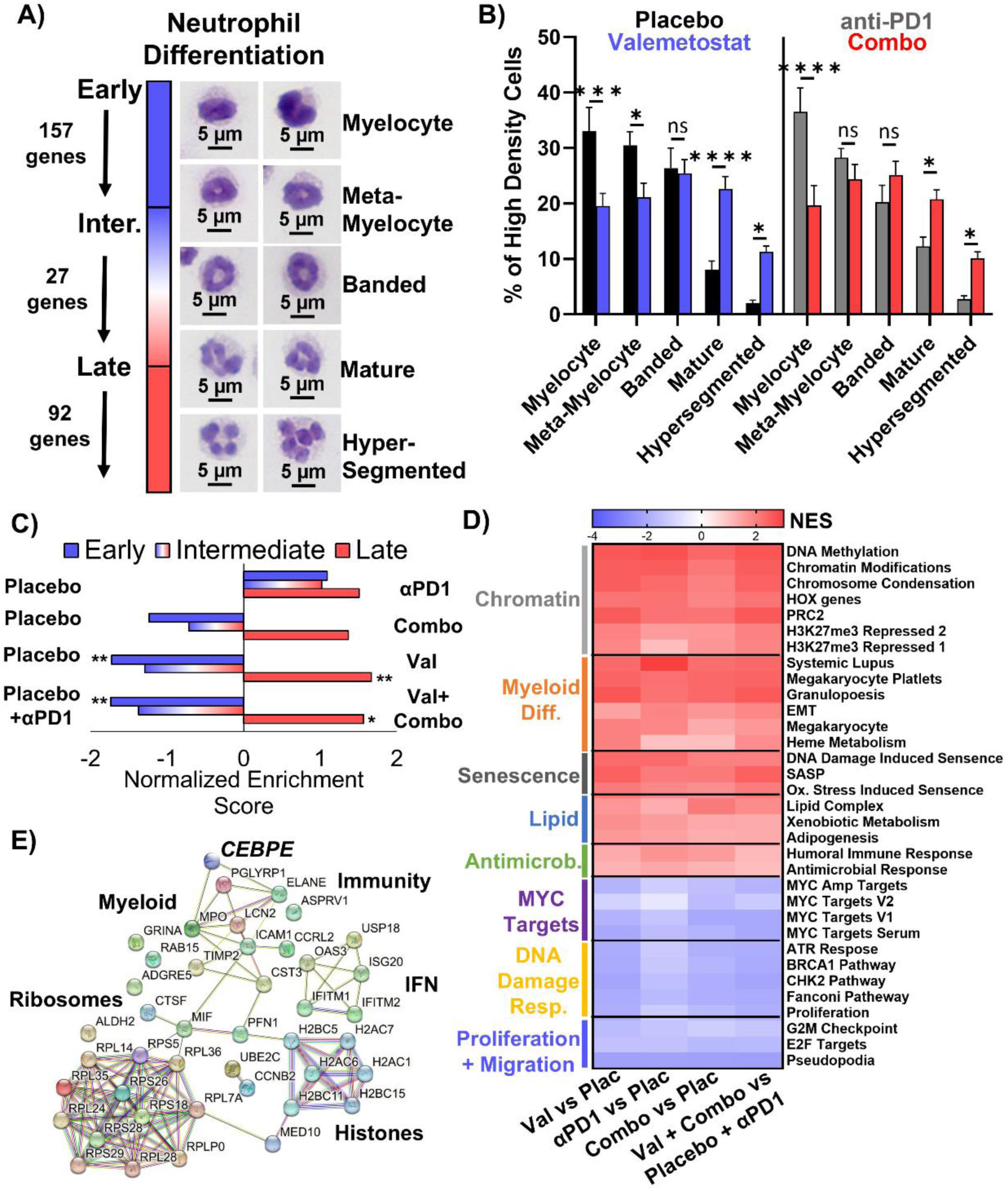
Neutrophils in bone marrow of valemetostat treated mice are more mature. **A)** Cytospin images of neutrophils in differing stages of differentiation. Representative images of nuclei types shown, scale bars = 5µm. **B)** Proportions of different nuclear morphologies in bone marrow cytospins, at least 71 neutrophils counted per sample, n=5 fpr placebo and combo, n=6 for valemetostat and anti-PD1, * indicates p<0.05 by two-way ANOVA with pairwise comparisons. **C)** Graph of normalized enriched scores for gene sets that correspond to three differentiation stages of neutrophils using indicated group contrasts. **D)** Heatmap of normalized enrichment score for indicated gene signatures using indicated group contrasts. **E**) StringDB analysis of mouse that are increased in valemetostat treated neutrophils See also Supp. Figure 3.

To visually represent the functional connectivity among genes involved in neutrophil differentiation, we utilized STRINGdb^29^, converting mouse genes to their human orthologs for this analysis. This network analysis highlights the complex interplay among genes driving neutrophil maturation (**Figure 3E, Supp. Table 2**). Notably, H2 histone variants were increased in the valemetostat treatment groups. We also observed upregulation of immune-related genes, including *CD80* and *B2M*, as well as genes associated with ribosomal function. *CEBPE*, a key transcriptional regulator of neutrophil maturation^30^, was also upregulated in response to treatment. Collectively, these data underscore the interconnected and complex transcriptional programs regulating neutrophil differentiation and highlight how EZH2 inhibition reshapes neutrophil identity toward a more mature, tumor-inhibitory state.

To further investigate the effects of *Ezh2* depletion on neutrophils, we utilized an *Ezh2* conditional deletion model (**Figure 4A, Supp. Figure 4A**). This model enables biallelic deletion of *Ezh2* upon administration of doxycycline for 7 days in the drinking water. Due doxycycline treatment of mice resulting in a depletion of the lower band, high density neutrophils, we implemented a three-day break following the treatment to allow for neutrophils numbers to recover (**Supp. Figure 4B**). To confirm doxycycline uptake across all mice, we assessed expression of the reporter gene H2B-GFP and observed slightly increased expression at 24 hours and modestly decreased expression at 48 hours in *Ezh2*-deleted (floxed) neutrophils (**Figure 4B**). RNA from these neutrophils was isolated, displaying decrease *Ezh2* expression with no changes in *Cxcr2* and *Mpo* (**Supp. Figure 4C**). We next evaluated neutrophil viability by assessing Annexin V staining and found that neutrophils with *Ezh2* deletion exhibited increased apoptosis at both 24 and 48 hours compared to controls. To determine whether EZH2 depletion affected neutrophil function, we performed an under-agarose migration assay in which neutrophils migrated toward a leukotriene B4 gradient. EZH2-deficient neutrophils demonstrated reduced migratory capacity, as visualized using both brightfield and GFP imaging (**Figure 4C**). To assess effector function, neutrophils were stimulated with phorbol 12-myristate 13-acetate (PMA) to induce neutrophil extracellular trap (NET) formation (**Supp. Figure 4D**). EZH2-deleted neutrophils exhibited a reduced frequency of NET formation relative to controls (**Figure 4D**). Finally, we evaluated antibacterial function and found that neutrophils from both *Ezh2* deficient and *Ezh2* proficient mice displayed comparable bacterial killing capacity (**Figure 4E, Supp Figure 4E**). Together these data suggest that although neutrophils may have impaired migration capacity when *Ezh2* is deficient, they are still able to perform their major function of bacterial killing and have more mature phenotypes including shorter half-lives. Ultimately, this combination of phenotypes may be more beneficial than neutrophil depletion, which is often associated with increased risks of infection.

**Figure 4:**
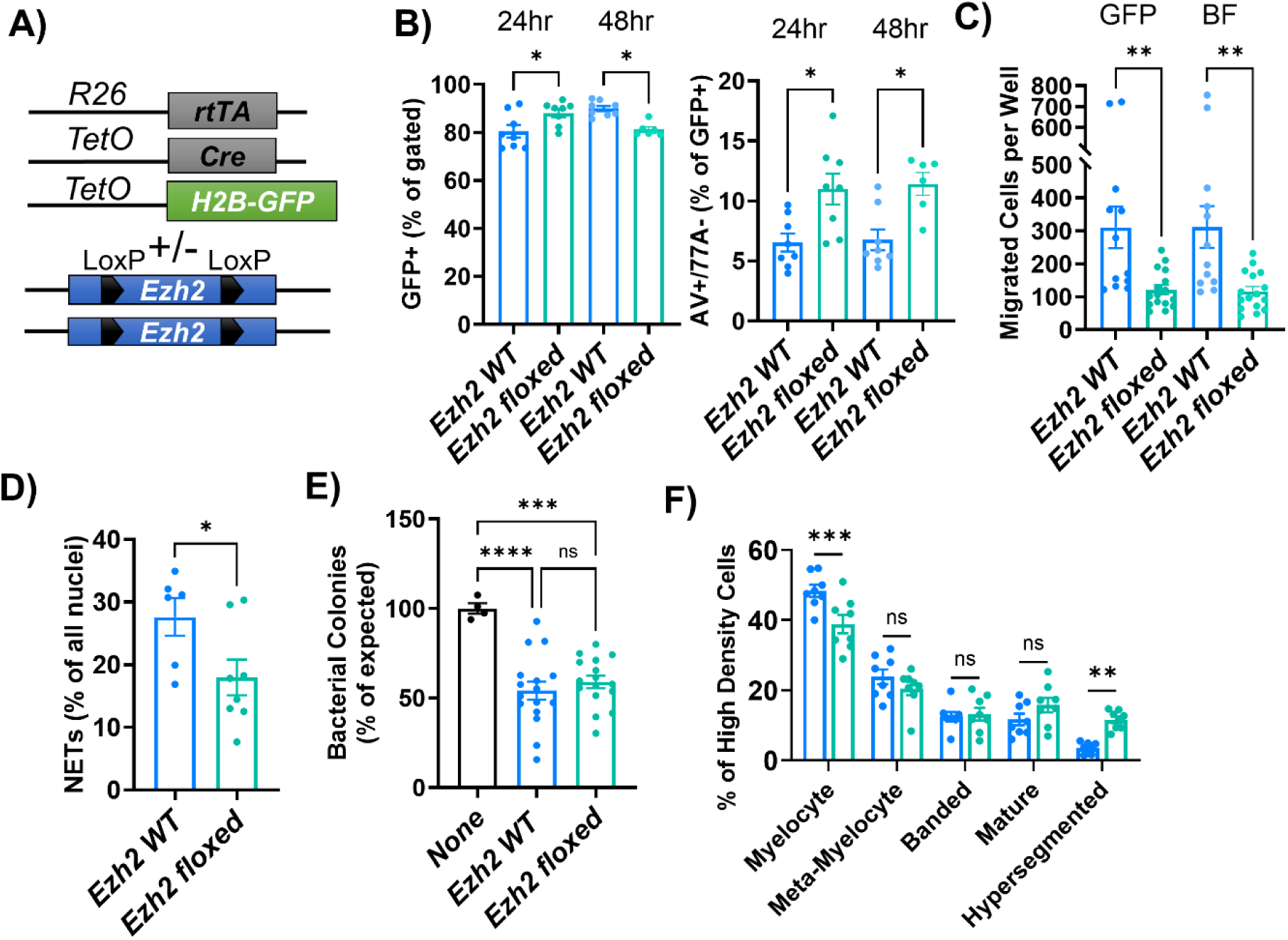
Functional assays of EZH2 depleted neutrophils. **A)** Allele schematic. **B)** At 24h and 48h, GFP tag and AV-apoptosis were quantified by flow cytometry, n=4 mice in duplicate per timepoint. **C)** At 18h, migration towards LTB4 was quantified by brightfield and fluorescence imaging, n=8 mice in duplicate. **D)** At 24h, percentage of NETs in response to PMA was quantified by fluorescence of GFP and DAPI, n=6 for WT and 8 for floxed. **E)** Neutrophils were incubated with E. coli and bacteria were plated and counted, n=8 mice in duplicate. **F)** Percentage of each nuclear morphology in cytospins of the high-density band, n=8 mice per genotype with at least 42 nuclei counter per mouse. For all graphs, mean +/- SEM is graphed, *p<0.05, **p<0.01, ***p<0.001, ****p<0.0001 by t-test **(B-D**), one-way ANOVA **(E)** or two-way ANOVA **(F**). See also Supp. Figure 4.

The tumor microenvironment (TME) is highly complex, and interactions among its cellular components critically influence therapeutic response and resistance. Modeling these interactions *in vitro* provides an opportunity to dissect mechanisms underlying treatment failure and immune evasion. To this end, we utilized three-dimensional tumoroid cultures that recapitulate the air-liquid-interface of the lung and mesenchymal interactions that we have used extensively for lung organoid and tumoroid growth. We first tested the ability of the lung mesenchymal cells to support tumoroid growth in a simplified media, and found that mesenchymal cells allowed for similar number of organoids to grow as when a complex “murine tracheal epithelial cell” media was used (**Supp. Figure 5A&B**). Using this base culture of tumoroids and mesenchymal cells in simplified media, we then added CD8+ T cells, and bone marrow (BM)–derived cells (**Figure 5A&B**). We chose bone marrow as the source of myeloid cells due to observing changes in response to valemetostat treatment, and due to the relatively immature status of bone marrow neutrophils relative to those in peripheral blood. This characteristic should allow bone marrow myeloid cells to persist in these long-term cultures. Bone marrow was isolated from two distinct conditions: tumor naïve mice, and mice harboring actively growing tumors. Notably, the addition of bone marrow from either condition resulted in a significant increase in tumoroid growth (**Figure 5C, Supp. Figure 5C**). This tumoroid growth-promoting effect of bone marrow was reversed in a stepwise fashion by treating the cultures with valemetostat, anti-PD1 antibody, or combination. We confirmed this was not due to general cell death in the cultures through CellTiter Glo analysis (**Figure 5D**). Flow cytometric analysis revealed a significantly higher number of CD8⁺ T cells in the combination treatment group at the conclusion of the experiment (**Figure 5E**). Among these CD8⁺ T cells, a greater proportion expressed PD-1 in the combination treatment compared to placebo (**Figure 5F&G**).

**Figure 5:**
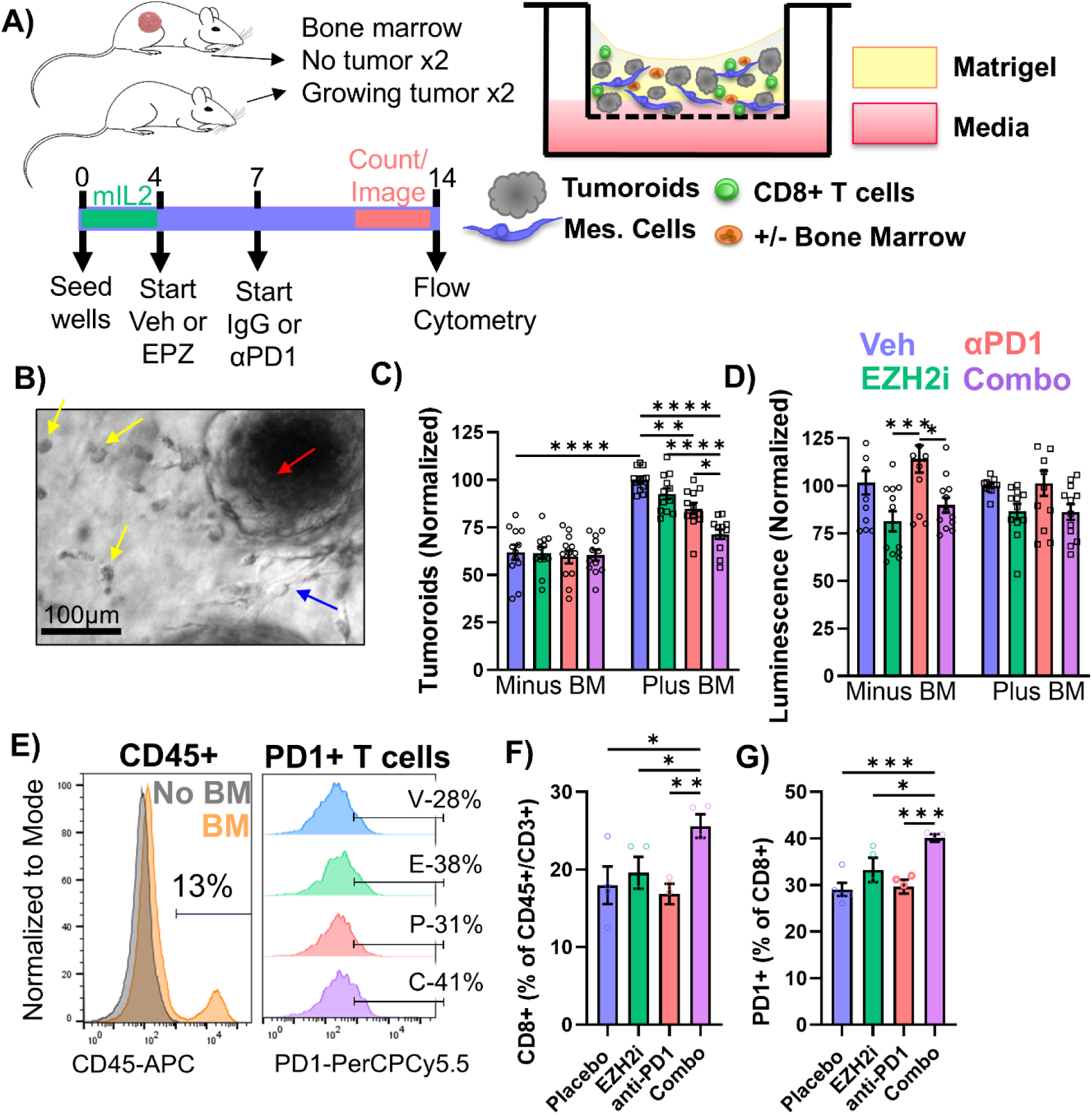
‘Multi-Cultures’ Recapitulate *in vivo* Tumor Responses. **A)** Schematic of air-liquid-interface tumoroid cultures with supporting lung mesenchymal cells and immune cells. **B)** Brightfield image of preliminary multi-culture, red arrow is *Lkb1/Pten* tumoroid, blue arrow is a group of lung endothelial cells, yellow arrows are small single cells that are likely bone-marrow derived. **C)** Tumoroid counts and normalize luminescence (CellTiter Glo) after indicated treatments, n=4 biological replicates for BM donors in 3 separate experiments, ***** p<0.0001, ***p=0.0006, ** p<0.01, *p<0.03 by two-way ANOVA with Holm-Šídák’s multiple comparisons test. **D)** Example of flow cytometry of multi-cultures that received indicated treatments. **E)** Flow cytometry for CD8+ cells as proportion of CD3+ in +BM cultures. **F)** Flow cytometry of PD1+ as proportion of CD8+ cells in +BM cultures. For all graphs mean +/- SEM is graphed, *p<0.05, **p<0.01, ***p<0.001, ****p<0.0001 by one-way ANOVA with pairwise comparisons. See also Supp. Figure 5

We next repeated these experiments using bone marrow derived from mice that had either spontaneously rejected tumors or from mice bearing actively growing tumors that had been treated with valemetostat (**Figure 6A&B**). Bone marrow from both tumor-rejected and valemetostat-treated mice inhibited tumoroid formation, with bone marrow from valemetostat-treated mice exhibiting the strongest inhibitory effect (**Figure 6C&D**). Flow cytometric analysis demonstrated a marked reduction in EpCAM+ tumor cells in cultures containing bone marrow from tumor-rejected or valemetostat-treated mice (**Figure 6E**). Furthermore, MHC class II expression was greatly increased in both rejected and valemetostat-treated bone marrow conditions relative to cultures without bone marrow (**Figure 6F, Supp. Figure 6A**). MHC class I and PD-L1 expression were significantly elevated in cultures containing tumor-rejected bone marrow, but were less increased in cultures receiving valemetostat-treated bone marrow (**Figure 6G, Supp. Figure 6B&C**). Examination of immune populations in these cultures revealed an increased proportions of CD45+ and CD8+ T cells in the valemetostat-treated bone marrow group, despite all conditions being seeded with equivalent numbers of T cells at baseline (**Figure 6H, Supp. Figure 6D**). Additionally, the valemetostat-treated bone marrow group exhibited an increased population of CD11b+ myeloid cells, and CD11b⁺Ly6G⁺ neutrophil (**Figure 6I, Supp. Figure 6E**). CellTiter Glo confirmed that despite the marked decrease in tumoroids, both the tumor-rejected and valemetostat-treated groups had slightly higher relative cell activity, likely from the increased immune cell populations (**Supp. Figure 6F**). We next carefully analyzed the bone marrow from each type of donor mouse used for the multi-culture experiments. By cytospin, we observed that valemetostat high-density neutrophils were more mature than tumor naïve or tumor-rejected (**Supp. Figure 6G**). Furthermore, when analyzing the exact bone marrow mixtures used for multi-cultures, it was evident that both valemetostat-treated and tumor-rejected bone marrow had more mature phenotypes than tumor-naïve or placebo-treated mice (**Supp. Figure 6H**). To investigate transcriptional changes underlying these phenotypic differences, we examined RNA sequencing data from rejected tumor bone marrow relative to the prior four treatment arm RNA sequencing. When comparing neutrophils from tumor-rejected mice to placebo treated tumor-bearing mice, RNA sequencing demonstrated enrichment of gene signatures associated with late-stage neutrophil differentiation, corroborating the cytospin findings (**Supp. Figure 6I**). Furthermore, genes associated with chromatin modification, myeloid differentiation, senescence, lipid metabolism, and antimicrobial defense were upregulated (**Figure 6J**), whereas MYC target genes, as well as pathways related to DNA damage and repair, proliferation, and migration, were downregulated. Integration of this dataset with neutrophils from valemetostat-treated mice (either single-agent or combination treatment) revealed transcriptional changes consistent with those observed *in vivo* (**Supp. Table 3**). Neutrophils derived from tumor-rejected bone marrow were enriched for all differentiation stages relative to placebo-treated mice (**Supp. Figure 6I**).

**Figure 6:**
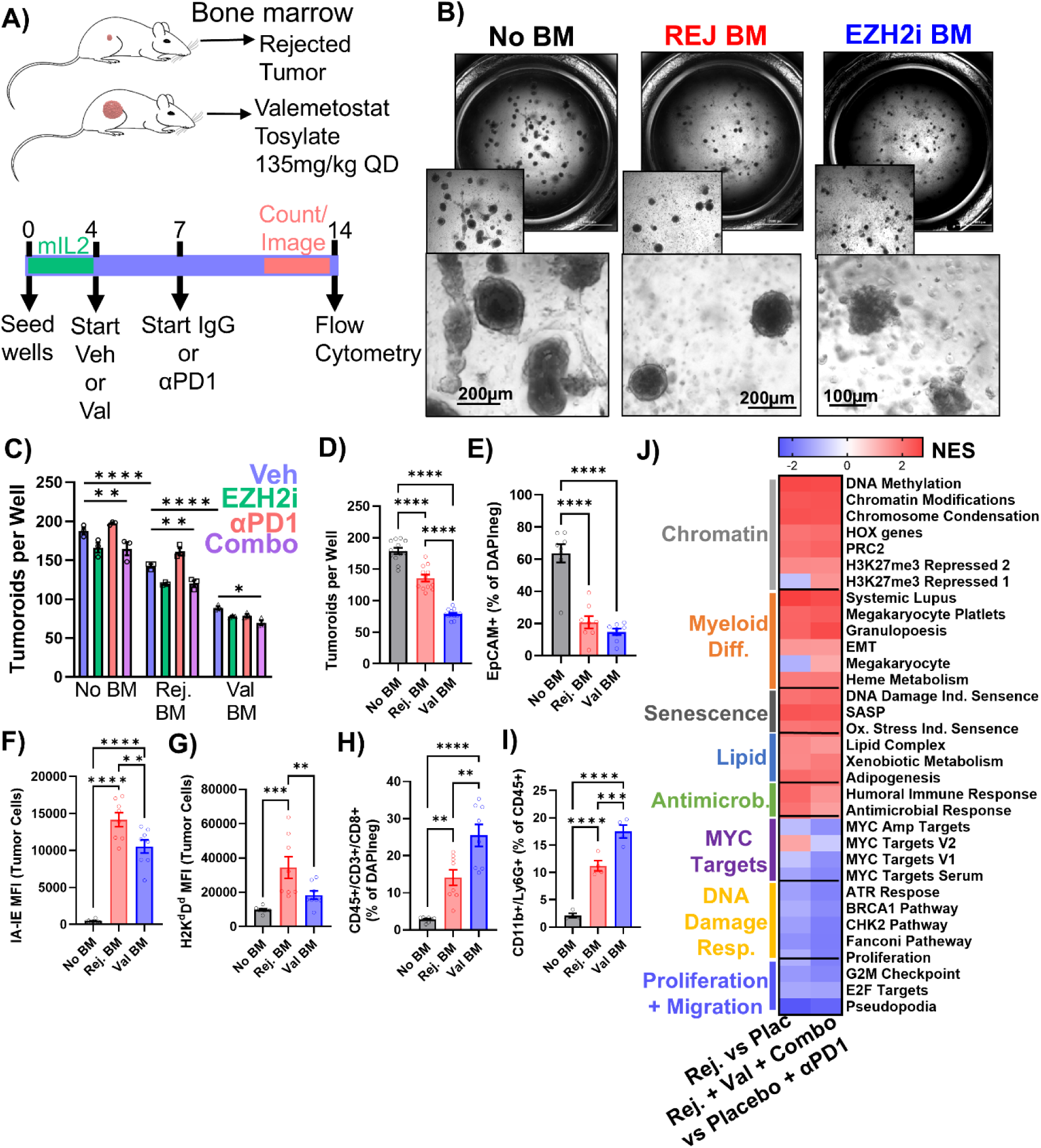
Bone marrow cells from valemetostat-treated or rejected tumor mice are anti-tumor. **A)** Schematic of samples used. **B)** Brightfield image of rejected bone marrow and valemetostat treated bone marrow cultures, zoomed scales = 200µm and 100µm. **C)** Tumoroids per well, n=3 wells per group, Tumoroids per well, n=3 wells per group, **** p<0.0001, **p<0.01, * p=0.04 by two-way ANOVA with Holm-Šídák’s multiple comparisons test. **D)** Tumoroids per well grouped by type of bone marrow, n=12 wells per condition. **E-J)** Flow cytometry n=8 wells from 2 separate experiments: **(E)** % EpCAM **(F)** MHC Class II IA-IE mean fluorescence intensity **(G)** MHC Class I H2K^d^D^d^ mean fluorescence intensity **(H)** % of CD45+/CD3+/CD8+, and **(I)** % of CD11b+/Ly6G+. For all graphs, mean +/- SEM is plotted, **** p<0.0001, *** p<0.001, ** p<0.01 by one-way ANOVA with pairwise comparisons. **J)** Heatmap of normalized enrichment score for indicated gene signatures using indicated group contrasts. See also Supp. Figure 6.

Given the consistent upregulation of MHC class II observed across both our *in vitro* and *in vivo* studies, we sought to determine the extent to which MHC II contributes to a full therapeutic response. To address this, we employed our multi-culture system using tumor-naïve bone marrow and total splenic T cells (**Figure 7A, Supp. Figure 7A**). Unexpectedly, blocking MHC class II abrogated the growth-promoting effect typically observed with the addition of bone marrow (**Figure 7B, Supp. Figure 7B**). Moreover, with MHC II blockade, none of the treatment conditions exhibited a reduction in tumoroid growth relative to placebo (**Figure 7C**). Flow cytometric analysis revealed no significant differences in the total number of CD45⁺ immune cells between MHC II–blocked and unblocked conditions (**Supp. Figure 7C**). While a trend toward reduced IFNγ expression in T cells was observed following MHC II blockade, this did not reach statistical significance (**Figure 7D**). However, MHC II blockade did result in a significant reduction in CD8⁺ T cell numbers in the presence of bone marrow (**Figure 7E**). MHC II blockade also altered antigen presentation on tumor cells – in the absence of bone marrow (-BM), MHC II blockade led to a marked decrease in MHC class I expression (**Figure 7F, Supp. Figure D**). In contrast, when bone marrow was present, MHC II blockade resulted in a substantial increase in MHC class I expression, and MHC II expression trended higher in the blockade group (**Supp. Figure 7E&F**). Additionally, a modest reduction in tumor cell PD-L1 expression was observed following MHC II blockade in the presence of bone marrow with a robust increase in without bone marrow present (**Figure 7G, Supp. Figure 7G**).

**Figure 7:**
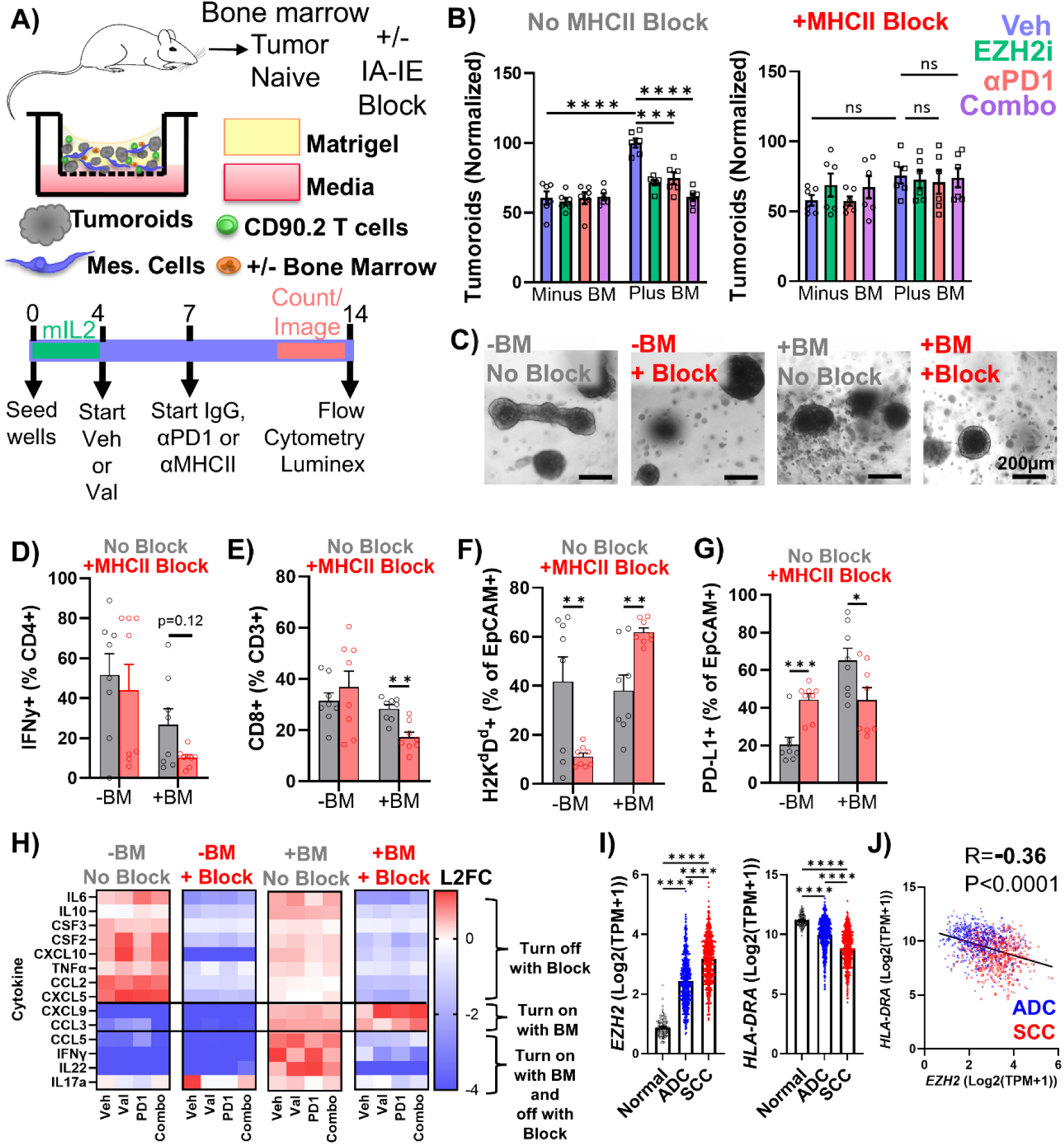
MHC Class II blockade limits bone marrow cytokine signaling. **A)** Schematic of samples used. **B)** Tumoroids per well, n=6 wells per group from two independent experiments, **** p<0.0001, ***p=0.0001 by two-way ANOVA with Holm-Šídák’s multiple comparisons test. **C)** Images of vehicle-treated wells of the indicated cultures are shown, scale bar = 200µm. **D-I)** Flow Cytometry **(D)** IFNγ as a % of CD4+ **(E)** CD8+ as a % of CD3+ **(F)** MHC class I H2KD + as a % of EpCAM+ **(C)** PD-L1+ as a % of EpCAM+. For all graphs, mean +/- SEM is plotted, ** p<0.01 and * p<0.05 by multiple un-paired t-tests between no block and block with Holm-Šídák’s *post hoc* correction. **H)** Log2 fold change in cytokine level in conditioned media relative to average for each cytokine. **I)** Relative expression of EZH2 in tumor-adjacent normal, lung adenocarcinoma (ADC) and lung squamous cell carcinoma (SCC), FDR for EZH2 ADC vs Normal 6.14e-69, SCC vs Normal 1.25e-126, ADC vs SCC 2.65e-42; FDR for HLA ADC vs Normal 6.19e-18, SCC vs Normal 5.44e-52, ADC vs SCC 4.54e-35. **J)** Pearson correlation coefficient of *EZH2* expression relative to *HLA-DRA* expression in ADC and SCC samples. See also Supp. Figure 7.

To further dissect the immunologic consequences of MHC II blockade, we performed cytokine profiling using a Luminex assay on culture supernatants collected at the conclusion of the experiment. In cultures lacking both bone marrow and MHC II blockade, multiple cytokines associated with tumor-promoting inflammation were detected, including IL-6, IL-10, CSF3, CSF2, CCL2, TNFα, and CXCL5 (**Figure 7H, Supp. Figure 7H**). Strikingly, the addition of the MHC II–blocking antibody in the absence of bone marrow eliminated the presence of these cytokines. When bone marrow was added without MHC II blockade, these cytokines re-emerged along with immune-activating mediators such as CXCL9, CCL3, CCL5, IFNγ, IL-22, and IL-17a. In contrast, cultures containing both bone marrow and MHC II blockade displayed a markedly restricted cytokine profile, with only CXCL9, CCL3, and CCL5 detected. These results suggest that MHC II blockade can broadly suppress both tumor-promoting inflammatory cytokines and immune-activating mediators, indicating that intact MHC II signaling may be required for both growth and elimination of tumor cells. Finally, to understand the prognostic value of these markers, we quantified the levels of *EZH2* and *HLA-DRA* (MHCII) transcripts in The Cancer Genome Atlas lung adenocarcinoma and lung squamous cell carcinoma datasets. We found that in lung squamous cell carcinoma, *EZH2* is highest and *HLA-DRA* is lowest when compared to normal adjacent lung or lung adenocarcinomas (**Figure 7I**). Furthermore, there is a strong negative correlation between *EZH2* and *HLA-DRA* in the malignant tissues (**Figure 7J**). Together, these data suggest that MHC class II signaling plays a central role in coordinating immune cell–derived cytokine networks within the tumor microenvironment, and may be specifically inducible in lung squamous cell carcinomas through EZH2 inhibition.

## Discussion

In this manuscript, we demonstrate that valemetostat, a dual EZH1/2 inhibitor, in combination with anti–PD-1 immunotherapy, effectively controls tumor growth in lung squamous cell carcinomas. This therapeutic effect is mediated by increased expression of MHC class I and class II, as well as PD-L1 on tumor cells, thereby enhancing antigen presentation and immune recognition. Simultaneously, valemetostat promotes neutrophil differentiation within the bone marrow and T cell activation in the tumor. RNA sequencing identified *CEBPE* as a gene upregulated in response to treatment. CEBPE is essential for proper neutrophil differentiation, as it regulates the formation of secondary and tertiary granules, and has been shown to suppress MYC target gene expression—consistent with our GSEA findings. Collectively, our data suggest that *CEBPE* may serve as a key driver of neutrophil differentiation in this context; however, further studies are required to determine whether this effect is a direct consequence of EZH2 inhibition or an indirect downstream outcome of treatment.

Through functional assays, we demonstrated that EZH2 deletion induces distinct changes in neutrophil behavior. Specifically, EZH2-deficient neutrophils exhibited a reduced capacity to form neutrophil extracellular traps (NETs), a process that has been linked to tumor metastasis. Consistent with this finding, our GSEA revealed upregulation of ALDH2, a known negative regulator of NET formation^31^. Functional assays also showed a decrease in neutrophil migratory capacity following EZH2 deletion, aligning with observations reported by other groups. Importantly, no differences were observed in bacterial killing capacity between wild-type and EZH2-deleted neutrophils. In agreement with these functional data, GSEA on valemetostat-treated neutrophils demonstrated enrichment of genes associated with antimicrobial responses, suggesting that EZH1/2-deficient neutrophils retain their ability to effectively combat infections. This result is consistent with no reports of increased bacterial infections in valemetostat treated patients.

Our multi-culture experiments successfully recapitulated key features of the *in vivo* tumor environment. These studies highlighted the critical importance of bone marrow origin, as naïve bone marrow or bone marrow from mice with actively growing tumors promoted tumoroid growth, whereas bone marrow from tumor-rejected or valemetostat-treated mice inhibited growth. These findings underscore the profound influence tumors exert on infiltrating immune cells and suggest that therapeutic reprogramming of these cells represents a viable strategy to improve treatment outcomes. Within the valemetostat-treated and tumor-rejected conditions, we identified neutrophil differentiation as a potential driver of tumoroid elimination. However, additional studies are required to determine whether neutrophils directly mediate tumor clearance, enhance T cell function, or act through a combination of both mechanisms. Notably, the observed tumoroid inhibition further suggests that tumors fully eliminated by this treatment strategy may be less likely to recur; however, this hypothesis will need to be formally tested in future studies.

Given the consistent upregulation of MHC class II observed throughout our studies, we sought to determine the importance of this axis in mediating treatment response. Blockade of MHC class II resulted in pronounced alterations in extracellular protein expression and cytokine profiles within the culture media. In the absence of bone marrow, MHC class II blockade led to a marked reduction in MHC class I expression that may result from the loss of CD8⁺ T cells that typically produce signals to increase MHC class I expression. In contrast, when bone marrow was present alongside MHC class II blockade, MHC class I expression was substantially increased, likely driven by bone marrow-derived immune cells enhancing tumor immunogenicity. Cytokine profiling further revealed that the presence of bone marrow—regardless of MHC class II blockade—was associated with expression of CXCL9 and CCL3, both of which are known to promote T cell recruitment and activation^32^. The observed increase in MHC class I expression may also be influenced by IL-22–mediated modulation of INFγ signaling. IL-22 is a known STAT3 agonist that suppresses INFγ-dependent gene expression^33^. In cultures containing bone marrow without MHC class II blockade, elevated IL-22 levels may suppress INFγ signaling, resulting in reduced MHC class I expression. Conversely, in the presence of MHC class II blockade, IL-22 levels are reduced, thereby permitting increased INFγ-driven gene expression and enhanced MHC class I upregulation. Further *in vivo* studies are required to validate these findings. Importantly, *in vivo* models will also allow us to determine whether the effects of MHC class II blockade can be overcome in the context of a fully intact immune system.

Collectively, these data reveal a previously underappreciated role for bone marrow–derived immune cells in driving tumor growth and therapeutic response. Specifically, our findings support a a model in which EZH1/2 inhibition promotes neutrophil maturation and functional reprogramming, generating bone marrow cells that restrain tumor growth and enhance anti-tumor immunity. Given that chemotherapy and tumors are known to drive ‘emergency myelopoiesis’ that results in a large production of immature neutrophils, it is tempting to speculate that EZH1/2 inhibition can reverse this phenomenon. These results underscore the importance of systemic immune remodeling in mediating durable responses to combination epigenetic and immunotherapy strategies.

## METHODS

### Mouse Experiments

All experimental animals were handled and cared for according to the University of Kentucky Institutional Animal Care and Use Committee (IACUC) guidelines. For syngeneic grafts, parental *Lkb1/Pten* floxed mice (FVB strain) were injected in each lower flank with a mixture of *Lkb1/Pten* tumoroids^11^ and Matrigel 1:1. Approximately 150,000 cells were seeded on each flank and allowed to grow for 6-8 weeks. These mice were then placed on the four treatment groups, placebo, anti-PD1, valemetostat, or combination treatment. Valemetostat tosylate provided by Daiichi Sankyo was prepared in sterile distilled water to a concentration 13.5mg/ml and administer via oral gavage at 135 mg/kg (equivalent to 100mg/kg valmetotstat). Anti-PD1 clone RMP1-14 (BioXCell #BP0146) and IgG2a isotype control (BioXCell #BP0089) were diluted with InVivoPure pH 7.0 dilution buffer (BioXCell #IP0070) or InVivoPure pH 6.5 dilution buffer (BioXCell # IP0065) to a concentration of 1.25μg/μL. Both antibodies were administered 7mg/kg 3 days a week via intraperitoneal injection. Tumors were measured via caliper where both length and width of each tumor were measured every two days during treatment. Tumor volume in mm^3^ was calculated by the formula length x width x width/2. Mouse health was monitored daily by weight and appearance judging the mice by: feel of the spine, coat appearance, alertness, and mobility. This score was recorded daily throughout the duration of the experiment.

The *Ezh2* conditional knockout model was produced previously^34^ by breeding Ezh2 floxed mice^35^ to mice containing *TetO:Cre*, *TetO:H2B-GFP* and *R26:rtTA*^34^. This mouse model allows for the biallelic deletion of EZH2 through the addition of 250mg of doxycycline (SIGMA, D9891-10G) and 5g of sucrose to 250mL water. Littermates that were either *Ezh2^flox/flox^* or *Ezh2^+/+^* with *TetO:H2B-GFP, Rosa26:rtTA and TetO:Cre.* EZH2 status as well as total genotype was confirmed on via genotyping prior to the experiment and deletion status was confirmed via genotyping. All mouse experiments were approved by the University of Kentucky Institutional Animal Care and Use Committees.

### Tumor Collection

Tumors were collected following mouse euthanasia. A portion of each tumor was placed in a tissue cassette, which was then placed into 10% formalin for at least 24 hours. Cassettes were moved to 70% ethanol and sent for paraffin embedding and sectioning by Markey Cancer Center Biospecimen Procurement and Translational Pathology Share Resource. 4µm serial sections were cut for each cassette. H&E-stained sections were analyzed using HALO nuclear profiler algorithm^24^. Additional sections were stained using Immunohistochemistry (IHC) using anti-H3K27me3 (1:500, Cell Signaling, C3B611) and anti-EZH2 (1:300, Cell signaling, #5246S) following our established protocols^34,36^. Staining was quantified in a blinded manner using the HALO software. The remaining portion of tumor was placed into a 1mL tube and minced using surgical scissors. 1mL of PBS and 20µl of collagenase dispase was added to each sample (Roche, 77425920), which was then rotated for 45 minutes at 37°C. After 45 minutes, tubes were put on ice, and 7.5µL of 3.7mg/mL of DNase I (SIGMA, D4527-10KU) was added. The digested tissue was then filtered first through a 100µm filter (VWR, 76327-102) then through a 40µm filter (VWR, 76327-098). The resulting cells were pelleted and resuspended for flow cytometry described below.

### Neutrophil Isolation and Assays

Bone marrow from femur and tibia of mice was flushed using 1mL of Mg^2+^ and Ca^2+^ free PBS (Cytiva cat# SH30256.01). 1.5 ml of histopaque 1077 was pipette on top of histopaque 1119 carefully to preserve the interface between the two layers. Total bone marrow (1mL) was then added on top of histopaque gradient and the tube was centrifuged at 1000x g for 25 minutes with zero acceleration and zero brake. Once centrifuged, the upper low-density and lower high-density bands of cells were collected and washed with excess PBS. Resulting cells were counted by hemacytometer and used within several hours for assays.

For cytopsin, neutrophils were resuspended in 150µL of Mg^2+^ and Ca^2+^ free PBS then spun onto a slide for 1 minute at 500 rpms. Wax circled spots were then fixed with 10% formalin for 15 minutes. Fixing agent was removed, and 150µL of PBS with 0.1% Triton-X was added for 15 minutes. The slides were then stained with Harris’s hematoxylin. Slides were moved back into organic solution through ethanol to xylenes. Cytoseal (Epredia, 8312-4) was used to afix the cover slips prior to imaging.

For neutrophil migration, six well plates were coated with FBS for 30 minutes. Once plates were coated, the gel mixture was created by combining 50mL of Mg^2+^ and Ca^2+^ free PBS and 0.6 grams of agarose (SIGMA, 9012-36-6). This was microwaved to mix the agarose with the PBS and then 50 mL of RPMI (Gibco, 11875-093) was added and mixed. 4.5mL of the mixture was pipetted into the six well plates that are then allowed to dry on a flat surface. Once wells were completely dry, a 3mm hole punch was used to create three holes exactly 2mm apart. Agarose plugs were then removed by aspiration, and the holes were washed with 1 ml of PBS. The plate was then put into the incubator and allowed to rest for 1 hour. Once the rest was completed the holes were aspirated and the chemo- attractant and control well were filled. The chemoattractant was 25µL of 250ng/mL Leukotriene B4 (Cayman, 20110) and the control was PBS. These were allowed 45 minutes to rest and create a gradient. 100,000 neutrophils (high-density band) were added to the center well and the outside wells are topped off. The neutrophils were placed in the incubator and allowed to migrate overnight. These were then fixed with formalin, and the wells were imaged on the Cytation5 (Biotek) using both bright field and GFP.

For the neutrophil extracellular trap (NET) assay, neutrophils were isolated following the above protocol. Neutrophils (high-density band) were then plated at 100,000 cells in 50µl of RPMI (Gibco, 11875-093) in chamber slides (Thermo Scientific, 154534). These were then placed in the incubator and allowed 30 minutes to rest. Once that was finished, Phorbol 12-myristate 13-acetate (Millipore, 524400) was diluted in RPMI at 4µL in 400µL of RPMI and 50µL of the dilution was added. These were allowed to sit for exactly 16 hours in the incubator. Once finished the chamber slide was removed and they were stained with DAPI prolonged gold (Invitrogen, P36935). The cover slip was added, and the slides were imaged using the Nikon TiE inverted microscope. These were quantified by visually counting for the NETs using DAPI and GFP fluorescence.

Isolated neutrophils (high density band) were plated at 10,000 cells per well in 100uL RPMI + 10% fetal bovine serum without antibiotic. The plate was allowed to rest in the 37°C, 5% CO_2_ incubator for 1 hour. Bacteria concentration (Stbl3 *E. coli* with pLK0.1 shGFP vector, Invitrogen) was measured using the OD_600_ of the bacteria grown in Luria broth with ampicillin. Triplicate OD_600_ values were averaged subtracting the average of the blank (no bacteria) wells. This number was then multiplied by eight to give the value. This value was diluted down to 0.1 in LB using *C*_1_*V*_1_ = *C*_2_*V*_2_ where *C*_1_ is the OD_600_ average, *V*_1_ is the unknown volume in µl, *C*_2_ is the target OD_600_ which for this case is 0.1, and *V*_2_ is the final volume of 1000µl. A new plate was created using the new diluted bacteria with each well in the 96 well plate still having a final volume of 200µL. The OD_600_ was measured run to ensure the 0.1 value was obtained. Vials with known concentration were frozen in 1:1 glycerol to bacteria suspension and stored at -80°C. After thawing, the number of bacteria needed to do a 1:1 incubation with 10,000 neutrophils was calculated, added to the 96 well plates with neutrophils and incubated together in 37°C 5% CO_2_ for 50 minutes. Total volumes of each well were collected and wells were washed with 200µL of PBS. Bacteria:neutrophil mixtures were pelleted and resuspended in 500µL of PBS. The number of desired colonies was calculated (10,000 bacteria in 500µl would require 25µl for 500 colonies). This amount was added to standard ampicillin agar plates and spread using sterile beads. Plates were incubated at 37°C for at least 24 hours. The plates were then removed and imaged. These images were then analyzed using the LifeScience Vision works software. This was manually checked to eliminate computer-based errors.

### T-Cell isolation

T cells were all isolated from the spleens of tumor bearing mice prior to addition into the multi-culture. They were also syngeneic with the tumor that was being used in the study. T cells were isolated using the Dynabeads FlowComp mouse CD8 kit (Invitrogen, 11462D) or CD90.2 (Invitrogen, 11465D) following the manufacture’s protocol. T cells that were frozen prior to multi-culture were stimulated prior to addition to the culture using 96 well plates with 2µg/mL CD3e (Invitrogen, 16-0031-82) and 5µg/mL CD28 antibodies (Invitrogen, 16-0281-82) precoated for 2 hours at 37% CO2 with the antibodies.

### Tumor Cell 3D cultures

Tumoroids previously isolated from our mouse models^11,25^ and seeded in growth factor reduced and phenol red-free Matrigel (Corning, 47743-722) in transwells with 0.4µm pore size (Corning 3470). Tumoroids were fed with DMEM/F12 media (ThermoFisher, 35050061) with 5µg/mL ITS (SIGMA, I3146), and 8-9% fetal bovine serum (VWR, 97068-085), termed ‘simple 3D media’. For murine tracheal epithelial cell media (MTEC)^37^, simple 3D media was supplemented with: 10µg/mL insulin (SIGMA, I0516), 0.1µg/mL cholera toxin (SIGMA, C8052), 25ng/mL EGF (Invitrogen, 53003-018), 25ng/mL bFGF (VWR, 10002-562) and 30µg/mL bovine pituitary extract (Invitrogen, 13028-014). Tumoroids actively growing in MTEC media were collected and trypsinized to single cell suspension for ‘multi-cultures’. Once combined with other cell types during the multi-culture experiments, additional supplements were not added and ‘simple 3D’ media was used. Lung mesenchymal cells from young C57bl/6 mice were isolated and cultured as previously described^34^.

The initial cultures contained cells that were combined at the following ratios: 100,000 neutrophils (combined both high and low density), 10-12,000 CD8+ T cells, 2,500-5,000 tumoroids, and 25,000 MECs. For the cultures originating from tumor rejected or valemetostat treated bone marrow, the cells were combined at the following rations: 100,000 neutrophils (70% low density and 30% high density), 4,000-5,000 CD8+ T cells, 5,000 tumor cells and 25,000 MECs. For the MHC block cultures, the following ratios were used: 5,000 tumor cells, 25,000 MECs, 100,0000 neutrophils (combined both low and high density), and T cells (CD90.2) at 10-20,000 per 200µL of 1:1 media to Matrigel mixture. Each treatment had three wells per experimental replicate. Cultures were allowed to establish in 100ng/mL mIL2 (Sino Biologics, cat #51061-MNAE) for 4 days prior to treatment. Wells were then placed on the different treatment arms, DMSO as vehicle control with 10µg/mL rat IgG2a (BioCell, BP0089) (Veh), 1µm valemetostat with 10µg/mL rat IgG2a (BioCell, BP0089) (Val), DMSO with 10µg/mL anti-PD1 (BioXCell BP0146 or BE0146) (PD1) or the combination of valemetostat with anti-PD1 (Combo). The valemetostat or DMSO was started on day 4 and the anti-PD1 or IgG2a started on day 7. Experiments with MHC II block were put into two categories, with 5µg/ml daily of anti-mouse I-A/I-E (Biolegend, 107602) or without. This was started on the same day as anti-PD1 and control IgG antibody treatment. All cultures were maintained at 37°C with 5% CO_2_. Wells were imaged using the Cytation5 and the tumoroid number was counted by eye under a light microscope.

For multiplex cytokine analysis, conditioned media was collected from the lower well of the transwell after at least 24 hours of incubation with the culture, and snap frozen in liquid nitrogen. Upon freeze thaw, 25uL was examined by Procarta custom mouse cytokine panel (ThermoFisher Scientific, Carlsbad, CA) on a Luminex 2000 machine according to the manufacturer’s instructions. Standard curves for each cytokine were used to calculate concentrations in media from triplicate wells. For tumoroid collection, all wells were treated with dispase (Roche diagnostics #77425920, RRID:SCR_011037) for at least an hour at 37°C with 5% CO_2_. Cell suspensions were collected, washed in PBS then trypsinized with 250µL and then neutralized with media. A portion of each suspension was placed into replicate wells in 96 well opaque plates. Then 50µL of CellTiter-Glo was added to each well, and after a 10-minute incubation, luminescence readings at gain 200 were acquired using a Cytation5 plate reader. The remainder was used for flow cytometry described below.

### Flow Cytometry

If starting from a culture, 150µL of dispase (Roche diagnostics #77425920, RRID:SCR_011037) was added for at least 1 hour at 37°C with CO2 prior to isolation of the cells. If using a solid tumor, the digestion protocol in “Tumor Collection” was followed. The samples were then washed with PBS and resuspended in 300µL of PF10 (PBS+10% fetal bovine serum (VWR, 97068-085)) + 4’,6-diamidion-2-phenylindole (DAPI, Sigma #D9542-5MG, 1:250). The samples were then stained using a combination of different panels to investigate the different populations present within the multi-culture or solid tumors. The flow cytometry antibodies that we used were; CD49F-FITC (Invitrogen #11-04-95-82, RRID:AB_11150059), PD-L1-PE (BioLegned #12430, RRID:AB_2073556), IA/IE-PERCP (BioLegend #107626, RRID:AB_2191071), EpCAM-PECy7 (BioLegend #18216, RRID:AB_1236471), CD31-APC (BioLegend #102510, RRID: AB_312917), Sca1-APCCy7 (BioLegend #108126, RRID:AB_10645327), CD8-FITC (BioLegend #100706, RRID:AB_312745), CD3-PE (BioLegend #100308, RRID:AB_312673), PD-1-PerCP (BioLegend #135208, RRID:AB_2159184), CD4-PECy7 (BioLegend #100528, RRID:312729), CD45-APC (BioLegend #103112, RRID:AB_312977), Ly6G-PerCP (BioLegend #127616 RRID:AB_1877271), CD11b-PECy7 (BioLegend #101216, RRID:AB_312799), Ly6A/E-APC (Thermo Fisher #17-5981-81, RRID:AB_469486), CD3-PE (BioLegend #100348, RRID:AB_2564038), CD8a-APC/Cy7 (BioLegend #100714, RRID:AB_312753), CD11c-PE (BioLegend #117308, RRID:AB_313777), H2Kd/H2Dd-AlexaF 647(BioLegend #114712, RRID:AB_493063), I-A/I-E-FITC (BioLegend #107605, RRID:AB_313320), CD81-PerCP/Cy5.5 (BioLegend #104911, RRID:AB_2562994), CD68-FITC (BioLegend #137005, RRID:AB_10575475), Ly6G-APC/cy7 (BioLegend #127623, RRID:AB_10645331), CD103-APC/Cy7 (BioLegend #121431, RRID:AB_2566551), CD63-PerCP/Cy5.5 (BioLegend #143911, RRID:AB_2565501), IFN-gamma-APC/Cy7 (BioLegend #505850, RRID:AB_2616698). For intracellular IFNγ, permeabilization buffer (Invitrogen, 00-8333-56) was used. The cells were incubated with the buffer for 5 minutes, then stained with IFNγ. After final wash, cells were fixed with 10% buffered formalin for 20 minutes, then moved into flow tubes through 35µm mesh caps. Data acquisition was performed on a BD Symphony2. For apoptosis assays, total neutrophils were plated at 100,000 cells per well in 96 well plates in media and run at 24 and 48 hours using Annexin V Apoptosis detection Kit (BioLegend, 640914). When experiments were conducted where the neutrophils were GFP+, the Annexin V PE (BioLegend, 640908) was used. All flow cytometry experiments were run using the BD FACSymphony (BioSciences) and analyzed using FlowJo.

### RNA isolation, RT-qPCR and Sequencing

All RNA isolations were done using Absolutely RNA Miniprep kits (Agilent, 400805) with DNase step using manufacturer’s instructions, and stored at -80°C until use. cDNA synthesis was performed using random hexamers and Super Script III (Invitrogen, 56575) using the manufacturer’s protocol, and RNase H (Ambion, AM2293) was added after reverse transcription to remove RNA template. cDNA was then diluted at 1:5 using nuclease free water and either stored at -80°C for later use or used to perform qPCR experiments. RT-qPCR master mixes were made for each gene of interest using the TaqMan fast advanced master mix (Invitrogen, 4444964) and TaqMan assays were run using the QuantStudio 3 (Applied Biosystems, A28567). All controls were run using *Gapdh* (Applied Biosystems, 4351309). Due to low RNA content in neutrophils, and testing for a gene that was knocked out, a maximum Ct value of 40 or 45 (depending on cycle used for run) was input when there were no Ct reads from any wells for that sample in that run.

RNAseq was performed by Innomics using mouse rRNA removal for library preparation and 150bp paired-end reads on a DNBseq platform. Data were aligned to mm38 using RSEM^38^, and edgeR^39^ was used to test expression differences. Log2-fold change of TPM counts from averages were used for heat maps in Supp. Tables. Due to yield, RNA used for RNA sequencing originated from the upper, low-density band of neutrophil isolation and at least two samples from each treatment group and sex were combined. No obvious sex differences were observed between groups, thus for total comparisons all groups we analyzed according to treatment. Gene set enrichment analysis^28^ using GSEA 4.3.2 was performed using log2-fold change ranked gene lists and the mouse to human ortholog mapping function. Hallmarks, GO-BP and C2 signatures were queried and selected enriched signature groups are shown. Neutrophil differentiation signatures were compiled from scRNAseq from three publications, including our own. StringDB^29^ was used to visualize networks up up-regulated genes. The data are available on Gene Expression Omnibus under record GSE330733.

### Patient Data Analysis

For lung cancer incidence, rates of age-adjusted invasive lung and bronchus tumors between 2013-2022 were extracted from National Program of Cancer Registries and Surveillance, Epidemiology, and End Results Program SEER*Stat Database: USCS Incidence Analytic Database with single ages and 90+ - 1998-2022 - linked to county attributes, United States Department of Health and Human Services, Centers for Disease Control and Prevention. Data were released in June 2025, based on the 2024 submission. The heatmap was composed of Q1-Q4 quartiles of the observed values by county level and NA were suppressed data. For lung cancer subtype distributions and survival, the following histology codes were used: Adenocarcinoma: 8140, 8253, 8254, 8255, 8260, 8265, 8480, 8481, 8490, 8551, 8570; Squamous Cell Carcinoma: 8070, 8071, 8072; Mixed and Not Otherwise Specified: 8000, 8010, 8022, 8046, 8230, 8560. Age-adjusted incidence rates and overall survival were extracted for invasive non-small cell lung cancers diagnosed between 2017 and 2022. Source data were from Surveillance, Epidemiology, and End Results (SEER) Research Plus Limited-Field Data, 22 Registries (Excluding IL and MA), Nov 2023 Sub (2000-2021), and Kentucky Cancer Registry (KCR) 2017-2021. Appalachian counties were used to segregate data as done previously^40^.

Gene-level RNA-seq expression and corresponding clinical data from The Cancer Genome Atlas (TCGA) for lung adenocarcinoma (ADC) and squamous cell carcinoma (SCC) were obtained from the Genomic Data Commons (GDC) using the GRCh38-based STAR counting workflow. To ensure a single RNA-seq profile per patient, technical and biological replicates were identified via TCGA barcodes. When multiple RNA-seq aliquots were available for a single patient, representative samples were selected using predefined analyte precedence rules (prioritizing RNA analytes in the order H > R > T) implemented in the IDConverter package. Remaining replicates were resolved based on portion and plate numbers using lexicographic ordering^41^ . Raw count and Transcripts Per Million (TPM) expression data from 109 normal tissues, 574 TCGA-LUAD and 552 TCGA-LUSC samples were retained for downstream analyses. Gene expression differences between tumor (ADC/SCC) and normal samples were evaluated using a linear mixed-effects model, with tissue type as a fixed effect and patient ID as a random intercept^42^. The independent ADC versus SCC comparison was assessed using a Mann-Whitney U test. Raw P-values across all three comparisons were pooled and adjusted by the Benjamini-Hochberg (FDR) method, with adjusted P < 0.05 considered significant. The correlation between gene expression was evaluated using a Pearson correlation test.

## ACKNOWLEDGEMENTS

This work was supported in part by R01 HL170193, R01 CA237643, P20 GM121327, American Cancer Society Grants 133123-RSG-19-081-01-TBG, and the Markey Research Women Strong Distinguished Researcher Award (CFB), UL1 TR001998 Center for Clinical and Translational Science pilot award (CFB), NCI T32 CA165990 (DRP and CMG), K99 CA303792 (DRP), Markey STRONG Scholars Program through the American Cancer Society IRG-19-140-31 (EMK). This research was also supported by the Biostatistics & Bioinformatics, Cancer Research Informatics, Oncogenomics, Biospecimen Procurement & Translational Pathology, Flow Cytometry & Immune Monitoring Shared Resources, and pilot grant funding from the University of Kentucky Markey Cancer Center through P30 CA177558.

## CRediT AUTHOR STATMENT

**Avery R. Childress:** Data Curation, Formal Analysis, Methodology, Investigation, Writing- Original Draft, Writing- Reviewing and Editing; **Dave-Preston Esoe:** Methodology, Investigation; **Xiulong Song:** Investigation; **Christain M. Gosser:** Investigation; **Yindan Lin:** Investigation; **Daniel R. Plaugher:** Investigation; **Tanner J. DuCote**: Methodology, Investigation; **Kassandra J. Naughton**: Investigation; **Erika M. Skaggs**: Methodology, Investigation; **Harrison Yang:** Investigation; **Ryan Goettl:** Formal Analysis, Investigation **Jinpeng Liu:** Formal Analysis, Investigation, Supervision; **Zhonglin Hao:** Conceptualization, Writing – Review and Editing; **Albert E. Fliss:** Resources; **Daisuke Honma:** Resources, Writing- Reviewing and Editing; **Todd Burus:** Formal Analysis, Visualization, Investigation; **Feitong Lei:** Formal Analysis, Investigation, **Bin Huang:** Formal Analysis, Investigation, Supervision; **Ellen J. Beswick:** Investigation, Writing- Reviewing and Editing; **Christine F. Brainson:** Conceptualization, Data Curation, Formal Analysis, Funding Acquisition, Visualization, Methodology, Investigation, Supervision, Writing- Original Draft, Writing- Reviewing and Editing.

---

**Supplementary Figure 1:**
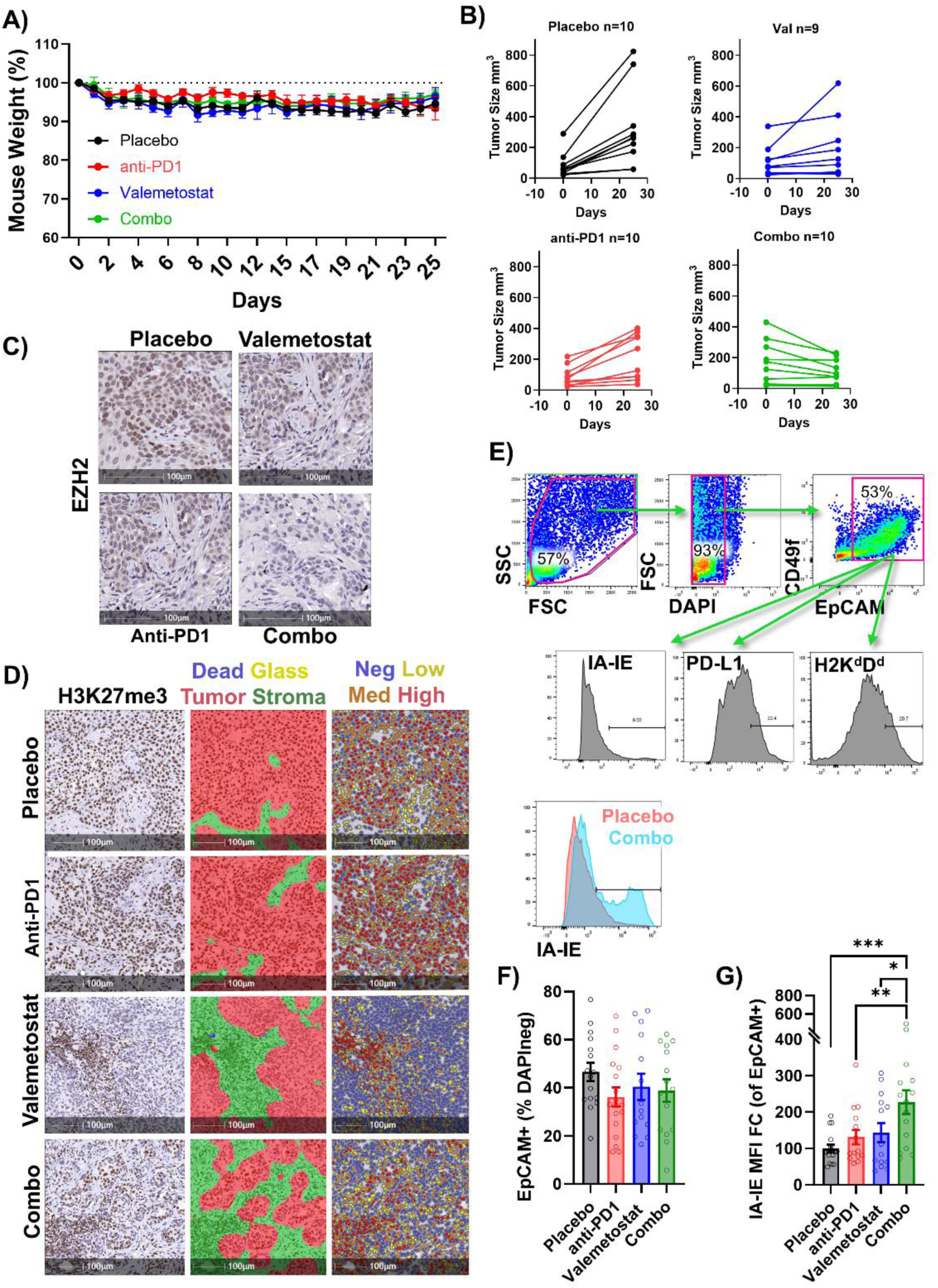
Related to Figure 1. **A)** Average weights of mice normalized to initial weight, mean +/- SEM is plotted. **B)** Initial and final tumor sizes (mm^3^) are plotted, n is indicated on graphs. **C)** Immunohistochemistry for EZH2, scale bars = 100µm. **D)** Flow cytometry for EpCAM+ cells as a percent of live gated cells in tumor samples mean +/- SEM is plotted. **E)** Representative flow cytometry gates for tumor samples with a histogram overlap of MHC Class II (IA-IE) in placebo vs a combination treated tumor. **F)** Immunohistochemistry for H3K27me3 with HALO tumor vs stroma and quantification algorithm, scale bars = 100µm. **G)** Gated on EpCAM+, IA-IE fold change with three outliers removed from anti-PD1 group by ROUT analysis. One-way ANOVA with pairwise comparisons shows significance.

**Supplementary Figure 2:**
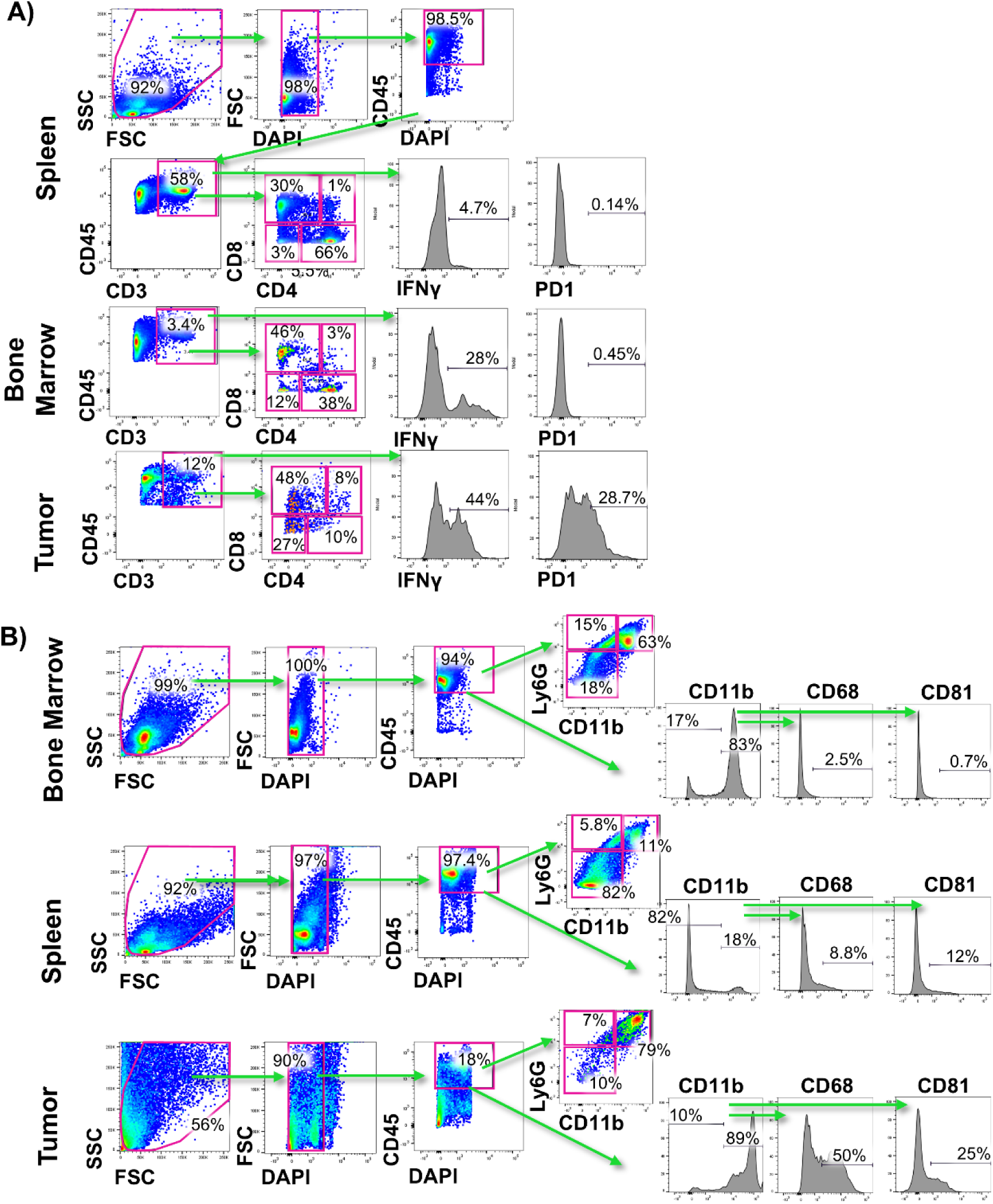
Related to Figure 2. **A)** Representative flow cytometry gating strategy for control immune cell population (spleens and bone marrows) and dissociated tumors from the syngeneic grafts from the indicated treatment for T cell stains. **B)** Representative flow cytometry gating strategy for control immune cell populations and dissociated tumors from the syngeneic grafts from the indicated treatment for myeloid cell stains.

**Supplementary Figure 3:**
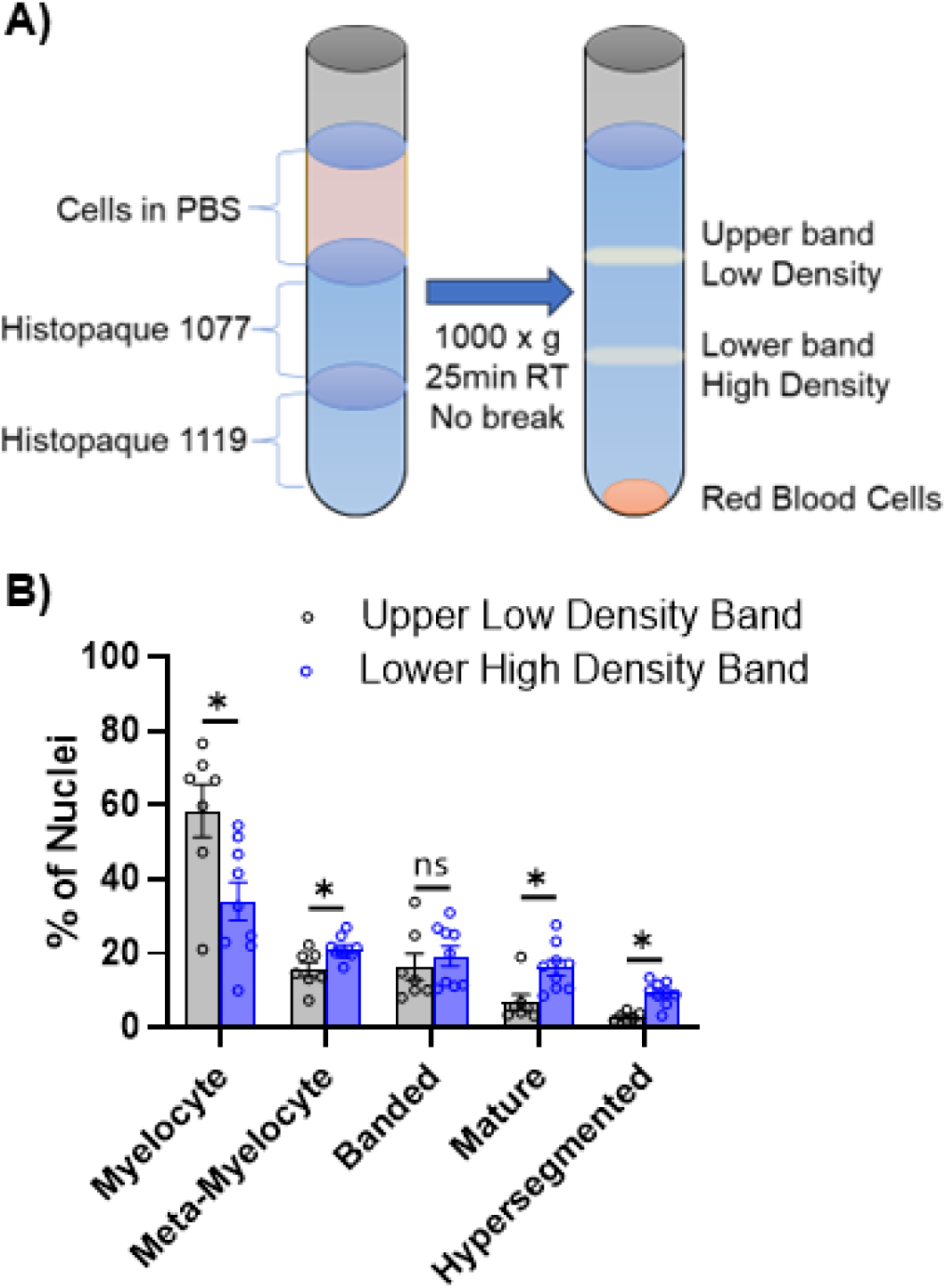
Related to Figure 3. **A)** Schematic of histopaque gradient used to isolate bone marrow cells and enrich for different cell types. **B)** Percentage of each nuclear morphology in cytospins of the high density band and low density bands, n=7 low density and n=9 high density samples with at least 42 nuclei counter per mouse, mean +/- SEM is plotted, * p<0.05 by multiple un-paired t-tests between high and low bands with Holm-Šídák’s *post hoc* correction.

**Supplementary Figure 4:**
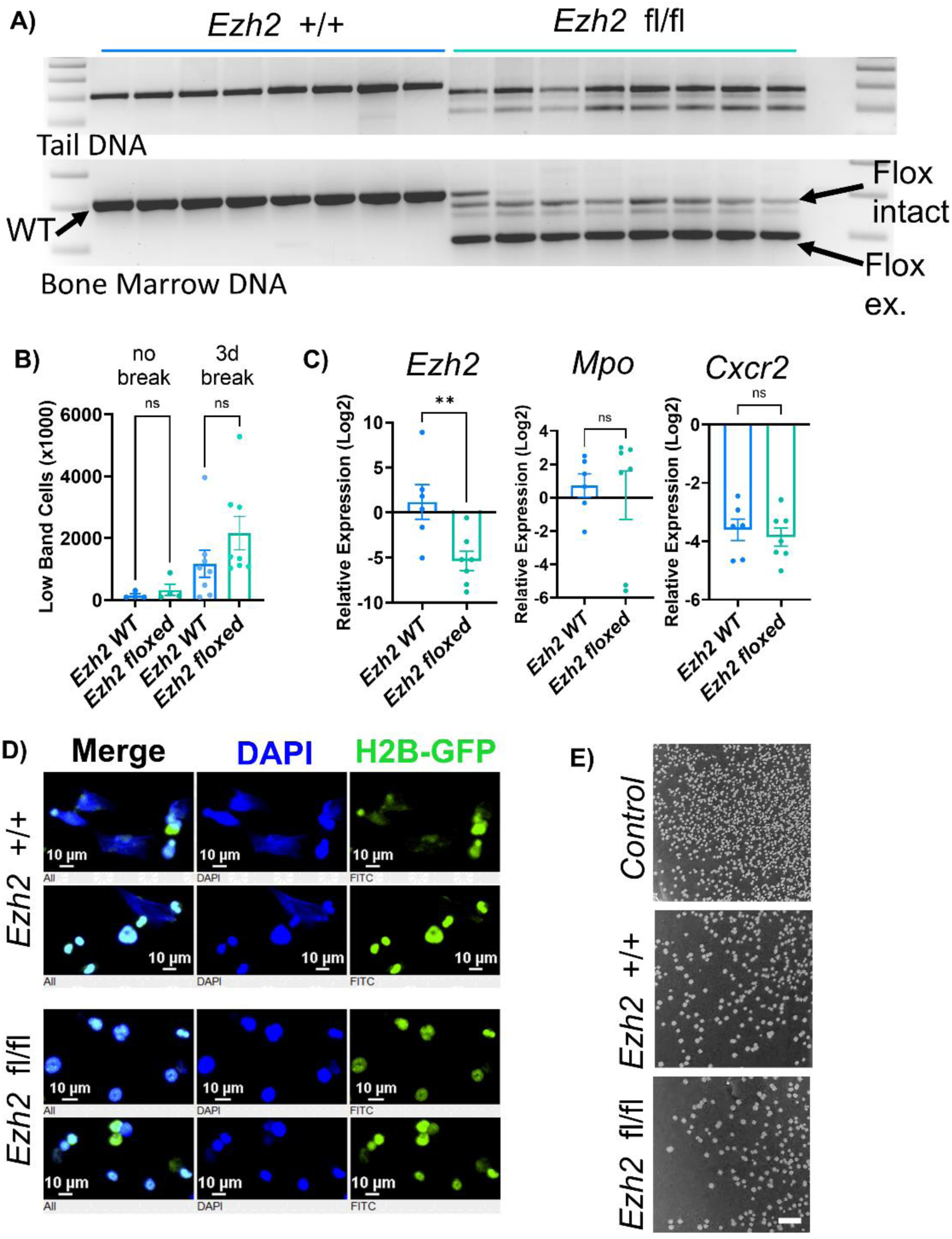
Related to Figure 4. **A)** Agarose gel showing DNA banding pattern after Ezh2 genotyping PCR on tail (top) and bone marrow (bottom). The first *Ezh*2 fl/fl sample was cross-contaminated with WT DNA and that sample was removed from other analyses of this population (RT-qPCR, apoptosis). **B)** Cells recovered from bone marrow extracted from both femurs and tibias with doxycycline in water until harvest date, or with doxycycline for 7 days then with a 3-day chase period prior to cell isolation. **C)** RT-qPCR of bone marrow (upper histopaque band) for *Ezh2*, *Mpo* and *Cxcr2*. Due to low RNA yield and testing for a gene that was knocked out, a maximum C_t_ value of 40 or 45 (depending on cycle used for run) was input when there were not Ct reads from any wells for that sample in that run. Four replicate plates were run from two different cDNA reactions and delta-Ct between multiplexed gene of interest and *Gapdh* were plotted. **D)** Representative images of PMA-induced NETs from bone marrow (lower histopaque band), scale bars = 10µm. **E)** Representative fields of bacteria from control (no neutrophils), WT or *Ezh2*-KO neutrophils, scale bars = 5mm.

**Supplementary Figure 5:**
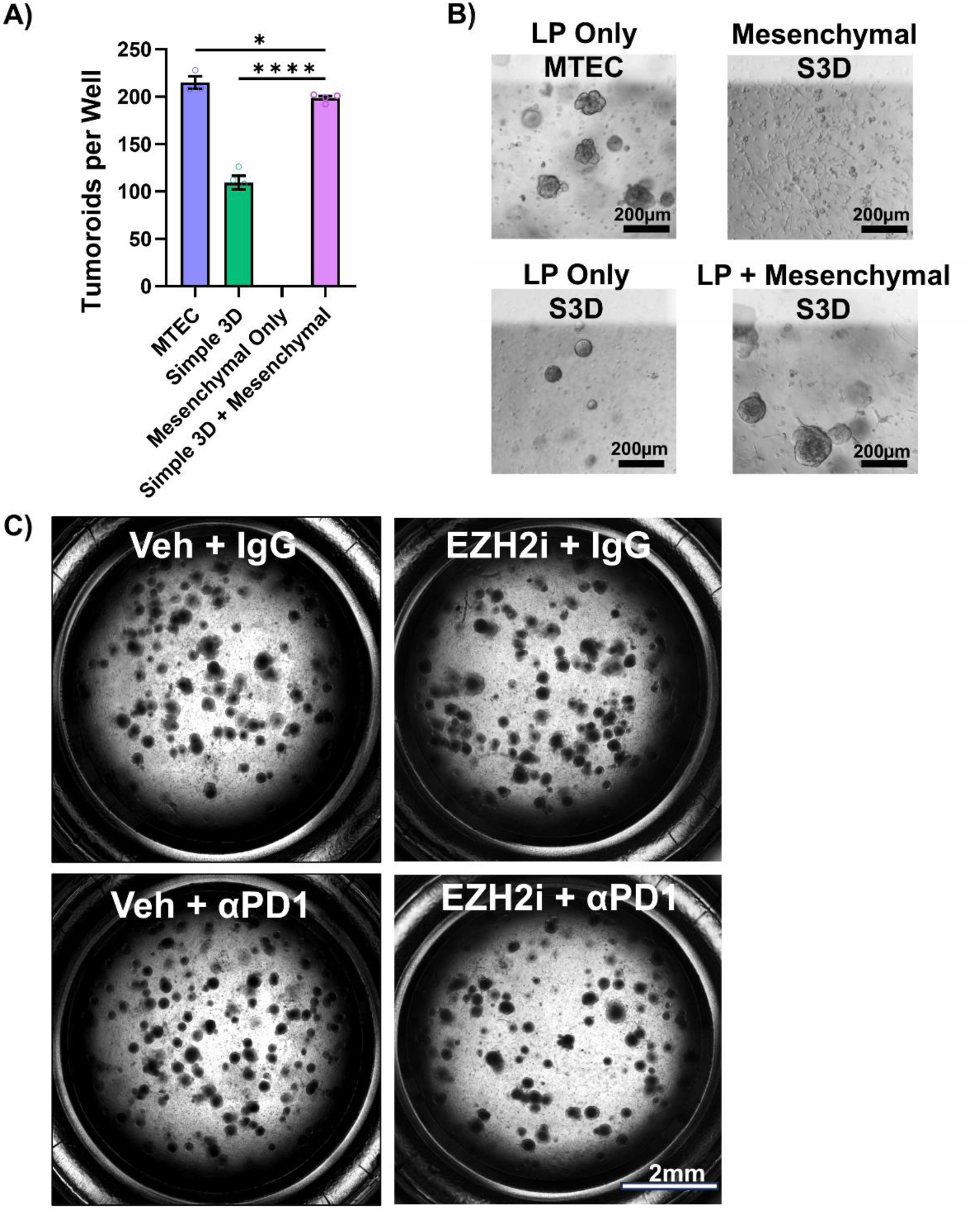
Related to Figure 5. **A)** Tumoroid counts in indicated culture conditions, n=3 wells for MTEC and 4 wells for other conditions. **B)** Brightfield images of indicated growth conditions, scale bar = 200µm. **C)** Brightfield images of bone marrow containing cultures treated with the indicated drugs, scale bar = 2mm

**Supplementary Figure 6:**
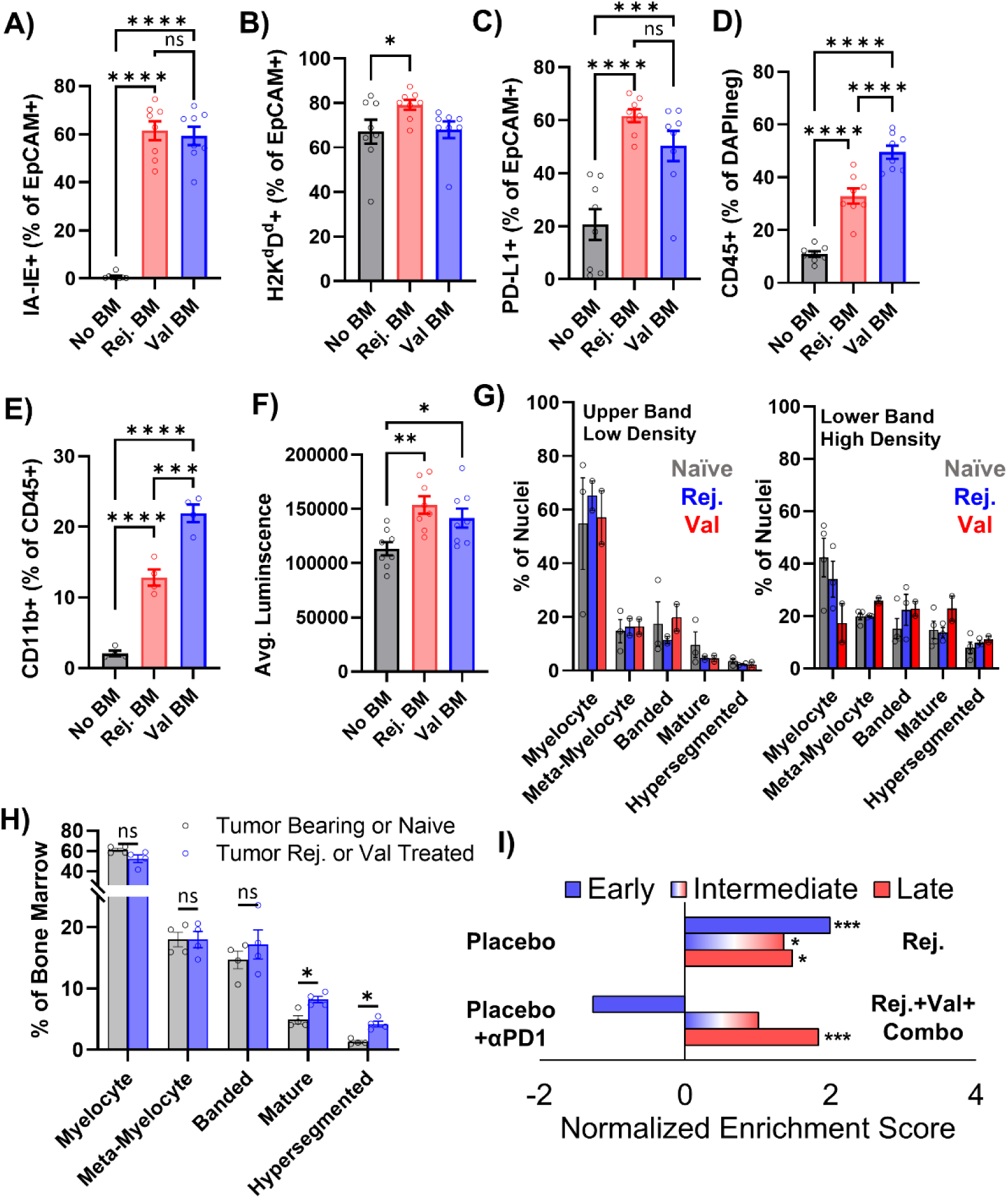
Related to Figure 6. **A-E)** Flow cytometry of **(A)** MHC Class II IA-IE % of EPCAM+ **(B)** MHC Class I H2K^d^D^d^ as a % of EPCAM+ **(C)** PD-L1+ as % of EPCAM+ **(D)** CD45 as a % of DAPI negative **(E)** CD11b+ % of CD45+. **F)** Average luminescence of each well measured by CellTiterGlo. **G+H)** Percentage of each nuclear morphology in cytospins of indicated density bands for samples used for multi-culture, **(G)** or bone marrow after remixing bands for multi-culture **(H)**. For all graphs, mean +/- SEM is plotted, **** p<0.0001, *** p<0.001, ** p<0.01 by one-way ANOVA with pairwise comparisons, except (H) which is multiple un-paired t-tests between groups with Holm-Šídák’s *post hoc* correction. **I)** Bar graph shows normalized enrichment score for three gene signatures for indicated contrasts.

**Supplementary Figure 7:**
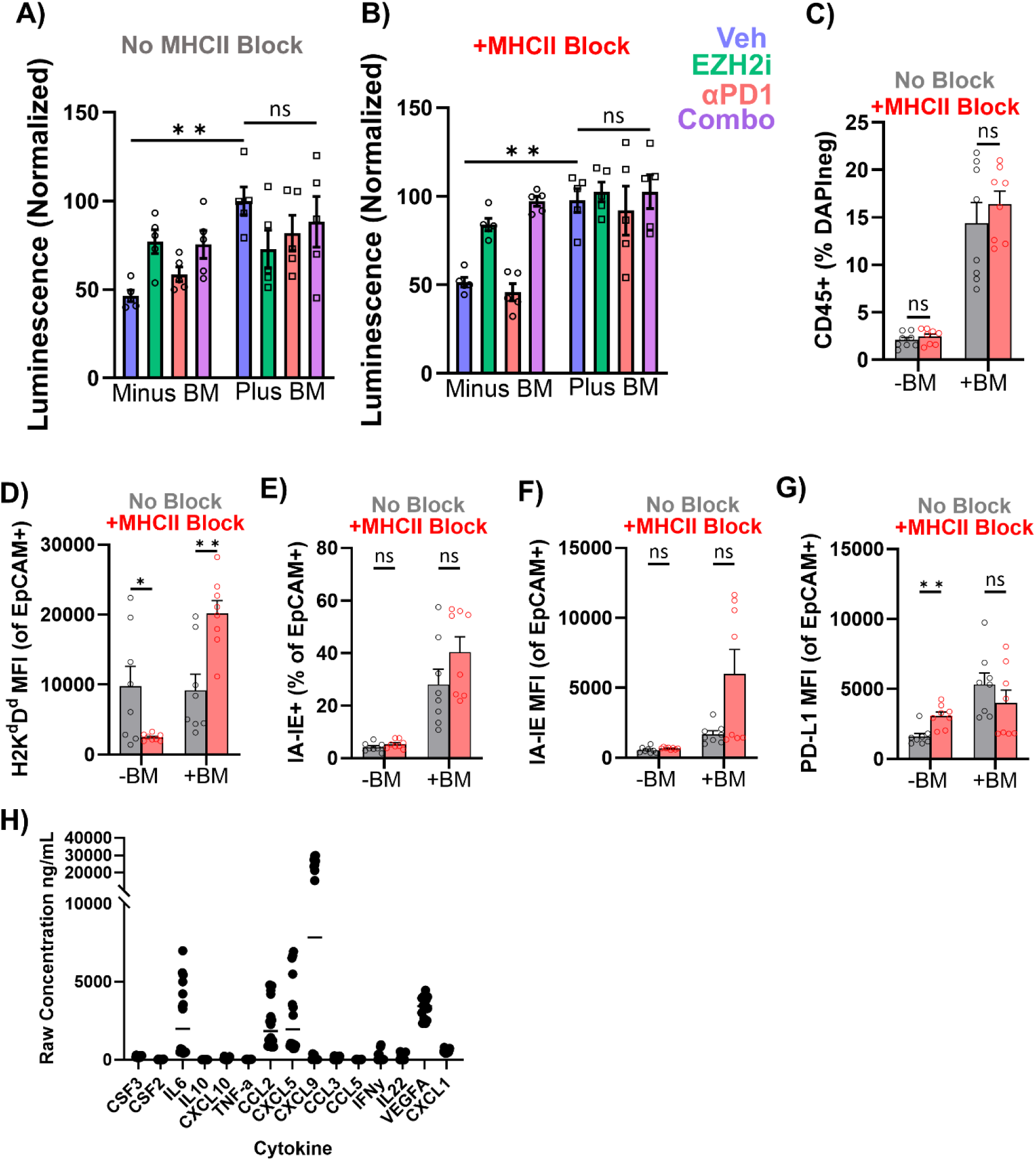
Related to Figure 7. **A)** CellTiter Glo n=2-3 wells per group from two independent experiments without MHC II block. **B)** CellTiter Glo n=2-3 wells per group from two independent experiments with MHC II block. Both A&B ** p<0.01 by two-way ANOVA with Holm-Šídák’s multiple comparisons test. **(C-G)** Flow Cytometry for: **(C)** CD45+ as a % of DAPI negative **(D)** MHC class I H2K^d^D^d^ normalized mean florescence intensity **(E)** MHC class II IA-IE as a % of EpCAM+ **(F)** MHC class II IA-IE mean florescence intensity. **(G)** PD-L1 normalized mean florescence intensity. **(H)** Concentration of cytokines in supernatants. For all graphs, mean +/- SEM is plotted, , ** p<0.01 and * p<0.05 by multiple un-paired t-tests between no block and block with Holm-Šídák’s *post hoc* correction

## REFERENCES

1. Borghaei, H., Gettinger, S., Vokes, E.E., Chow, L.Q.M., Burgio, M.A., de Castro Carpeno, J., et al. Five-Year Outcomes From the Randomized, Phase III Trials CheckMate 017 and 057: Nivolumab Versus Docetaxel in Previously Treated Non–Small-Cell Lung Cancer. Journal of Clinical Oncology 39, 723–733 (2021).

2. de Castro, G., Jr., Kudaba, I., Wu, Y.L., Lopes, G., Kowalski, D.M., Turna, H.Z., et al. Five-Year Outcomes With Pembrolizumab Versus Chemotherapy as First-Line Therapy in Patients With Non-Small-Cell Lung Cancer and Programmed Death Ligand-1 Tumor Proportion Score ≥ 1% in the KEYNOTE-042 Study. J Clin Oncol 41, 1986–1991 (2023).

3. Brahmer, J., Reckamp, K.L., Baas, P., Crinò, L., Eberhardt, W.E.E., Poddubskaya, E., et al. Nivolumab versus Docetaxel in Advanced Squamous-Cell Non–Small-Cell Lung Cancer. New England Journal of Medicine 373, 123–135 (2015).

4. Drilon, A., Rekhtman, N., Ladanyi, M. & Paik, P. Squamous-cell carcinomas of the lung: emerging biology, controversies, and the promise of targeted therapy. The Lancet Oncology 13, e418–e426 (2012).

5. Kargl, J., Busch, S.E., Yang, G.H.Y., Kim, K.-H., Hanke, M.L., Metz, H.E., et al. Neutrophils dominate the immune cell composition in non-small cell lung cancer. Nature Communications 8, 14381 (2017).

6. Salcher, S., Sturm, G., Horvath, L., Untergasser, G., Kuempers, C., Fotakis, G., et al. High-resolution single-cell atlas reveals diversity and plasticity of tissue-resident neutrophils in non-small cell lung cancer. Cancer Cell 40, 1503–1520.e1508 (2022).

7. Bracken, A.P. & Helin, K. Polycomb group proteins: navigators of lineage pathways led astray in cancer. Nature Reviews Cancer 9, 773 (2009).

8. Burr, M.L., Sparbier, C.E., Chan, K.L., Chan, Y.-C., Kersbergen, A., Lam, E.Y.N., et al. An Evolutionarily Conserved Function of Polycomb Silences the MHC Class I Antigen Presentation Pathway and Enables Immune Evasion in Cancer. Cancer Cell 36, 385–401.e388 (2019).

9. Ennishi, D., Takata, K., Beguelin, W., Duns, G., Mottok, A., Farinha, P., et al. Molecular and Genetic Characterization of MHC Deficiency Identifies EZH2 as Therapeutic Target for Enhancing Immune Recognition. Cancer Discov 9, 546–563 (2019).

10. Piunti, A., Meghani, K., Yu, Y., Robertson, A.G., Podojil, J.R., McLaughlin, K.A., et al. Immune activation is essential for the antitumor activity of EZH2 inhibition in urothelial carcinoma. 8, eabo8043 (2022).

11. DuCote, T.J., Song, X., Naughton, K.J., Chen, F., Plaugher, D.R., Childress, A.R., et al. EZH2 Inhibition Promotes Tumor Immunogenicity in Lung Squamous Cell Carcinomas. Cancer Res Commun 4, 388–403 (2024).

12. Hoy, S.M. Tazemetostat: First Approval. Drugs 80, 513–521 (2020).

13. Straining, R. & Eighmy, W. Tazemetostat: EZH2 Inhibitor. J Adv Pract Oncol 13, 158–163 (2022).

14. Keam, S.J. Valemetostat Tosilate: First Approval. Drugs 82, 1621–1627 (2022).

15. Zinzani, P.L., Izutsu, K., Mehta-Shah, N., Barta, S.K., Ishitsuka, K., Córdoba, R., et al. Valemetostat for patients with relapsed or refractory peripheral T-cell lymphoma (VALENTINE-PTCL01): a multicentre, open-label, single-arm, phase 2 study. The Lancet Oncology 25, 1602–1613 (2024).

16. Hasegawa, K., Sato, A., Tanimura, K., Uemasu, K., Hamakawa, Y., Fuseya, Y., et al. Fraction of MHCII and EpCAM expression characterizes distal lung epithelial cells for alveolar type 2 cell isolation. Respiratory research 18, 150 (2017).

17. Strunz, M., Simon, L.M., Ansari, M., Kathiriya, J.J., Angelidis, I., Mayr, C.H., et al. Alveolar regeneration through a Krt8+ transitional stem cell state that persists in human lung fibrosis. Nat Commun 11, 3559 (2020).

18. Jenkins, R.W., Aref, A.R., Lizotte, P.H., Ivanova, E., Stinson, S., Zhou, C.W., et al. Ex Vivo Profiling of PD-1 Blockade Using Organotypic Tumor Spheroids. Cancer Discov 8, 196–215 (2018).

19. Neal, J.T., Li, X., Zhu, J., Giangarra, V., Grzeskowiak, C.L., Ju, J., et al. Organoid Modeling of the Tumor Immune Microenvironment. Cell 175, 1972–1988.e1916 (2018).

20. Lo, H.C., Choi, H., Kolahi, K., Rodriguez, S., Gonzalez, L., Chu, F., et al. A 3D tumor spheroid model with robust T cell infiltration for evaluating immune cell engagers. iScience 28(2025).

21. Selli, M.E., Landmann, J.H., Arveseth, C. & Singh, N. Inducing T-cell dysfunction by chronic stimulation of CAR-engineered T-cells targeting cancer cells in suspension cultures. STAR protocols 4(2023).

22. Ren, Y., Lin, Z., Li, S., Zhang, R., Lai, Z., Jiang, H., et al. An in vitro co-culture model with CAR-T cells, antigen-presenting cells, and tumor cells to evaluate CAR-T cell-induced cytokine release syndrome. Front Cell Infect Microbiol 16, 1721114 (2026).

23. Bracken, A.P., Pasini, D., Capra, M., Prosperini, E., Colli, E. & Helin, K. EZH2 is downstream of the pRB-E2F pathway, essential for proliferation and amplified in cancer. Embo j 22, 5323–5335 (2003).

24. DuCote, T.J., Naughton, K.J., Skaggs, E.M., Bocklage, T.J., Allison, D.B. & Brainson, C.F. Using artificial intelligence to identify tumor microenvironment heterogeneity in non-small cell lung cancers. Laboratory Investigation, 100176 (2023).

25. Xu, C., Fillmore, Christine M., Koyama, S., Wu, H., Zhao, Y., Chen, Z., et al. Loss of Lkb1 and Pten Leads to Lung Squamous Cell Carcinoma with Elevated PD-L1 Expression. Cancer Cell 25, 590–604 (2014).

26. Zilionis, R., Engblom, C., Pfirschke, C., Savova, V., Zemmour, D., Saatcioglu, H.D., et al. Single-Cell Transcriptomics of Human and Mouse Lung Cancers Reveals Conserved Myeloid Populations across Individuals and Species. Immunity 50, 1317–1334.e1310 (2019).

27. Tang, K.H., Li, S., Khodadadi-Jamayran, A., Jen, J., Han, H., Guidry, K., et al. Combined Inhibition of SHP2 and CXCR1/2 Promotes Antitumor T-cell Response in NSCLC. Cancer Discovery 12, 47–61 (2022).

28. Subramanian, A., Tamayo, P., Mootha, V.K., Mukherjee, S., Ebert, B.L., Gillette, M.A., et al. Gene set enrichment analysis: A knowledge-based approach for interpreting genome-wide expression profiles. Proceedings of the National Academy of Sciences 102, 15545–15550 (2005).

29. Szklarczyk, D., Gable, A.L., Lyon, D., Junge, A., Wyder, S., Huerta-Cepas, J., et al. STRING v11: protein-protein association networks with increased coverage, supporting functional discovery in genome-wide experimental datasets. Nucleic acids research 47, D607–D613 (2019).

30. Serwas, N.K., Huemer, J., Dieckmann, R., Mejstrikova, E., Garncarz, W., Litzman, J., et al. CEBPE-Mutant Specific Granule Deficiency Correlates With Aberrant Granule Organization and Substantial Proteome Alterations in Neutrophils. Frontiers in immunology 9, 588 (2018).

31. Yang, K., Gao, R., Chen, H., Hu, J., Zhang, P., Wei, X., et al. Myocardial reperfusion injury exacerbation due to ALDH2 deficiency is mediated by neutrophil extracellular traps and prevented by leukotriene C4 inhibition. Eur Heart J 45, 1662–1680 (2024).

32. Kang, T.G., Park, H.J., Moon, J., Lee, J.H. & Ha, S.-J. Enriching CCL3 in the Tumor Microenvironment Facilitates T cell Responses and Improves the Efficacy of Anti-PD-1 Therapy. Immune network 21(2021).

33. Pennino, D., Bhavsar, P.K., Effner, R., Avitabile, S., Venn, P., Quaranta, M., et al. IL-22 suppresses IFN-γ-mediated lung inflammation in asthmatic patients. The Journal of allergy and clinical immunology 131, 562–570 (2013).

34. Byrd, A.L., Qu, X., Lukyanchuk, A., Liu, J., Chen, F., Naughton, K.J., et al. Dysregulated Polycomb Repressive Complex 2 contributes to chronic obstructive pulmonary disease by rewiring stem cell fate. Stem Cell Reports 18, 289–304 (2023).

35. Shen, X., Liu, Y., Hsu, Y.-J., Fujiwara, Y., Kim, J., Mao, X., et al. EZH1 Mediates Methylation on Histone H3 Lysine 27 and Complements EZH2 in Maintaining Stem Cell Identity and Executing Pluripotency. Molecular Cell 32, 491–502 (2008).

36. Chen, F., Byrd, A.L., Liu, J., Flight, R.M., DuCote, T.J., Naughton, K.J., et al. Polycomb deficiency drives a FOXP2-high aggressive state targetable by epigenetic inhibitors. Nature Communications 14, 336 (2023).

37. Rock, J.R., Onaitis, M.W., Rawlins, E.L., Lu, Y., Clark, C.P., Xue, Y., et al. Basal cells as stem cells of the mouse trachea and human airway epithelium. Proceedings of the National Academy of Sciences of the United States of America 106, 12771–12775 (2009).

38. Li, B. & Dewey, C.N. RSEM: accurate transcript quantification from RNA-Seq data with or without a reference genome. BMC Bioinformatics 12, 323 (2011).

39. Robinson, M.D., McCarthy, D.J. & Smyth, G.K. edgeR: a Bioconductor package for differential expression analysis of digital gene expression data. Bioinformatics 26, 139–140 (2010).

40. Brainson, C.F., Huang, B., Chen, Q., McLouth, L.E., He, C., Hao, Z., et al. Description of a Lung Cancer Hotspot: Disparities in Lung Cancer Histology, Incidence, and Survival in Kentucky and Appalachian Kentucky. Clin Lung Cancer 22, e911–e920 (2021).

41. Wang, S., Li, H., Song, M., Tao, Z., Wu, T., He, Z., et al. Copy number signature analysis tool and its application in prostate cancer reveals distinct mutational processes and clinical outcomes. PLoS genetics 17, e1009557 (2021).

42. Bates, D., Mächler, M., Bolker, B. & Walker, S. Fitting Linear Mixed-Effects Models Using lme4. Journal of Statistical Software 67, 1–48 (2015).

