## Supplemental Tables 1-3 for "EZH1/2 inhibition improves immunotherapy response through MHC Class II de-repression and neutrophil reprogramming"

Childress et al.,

Supplemental Tables 1-3

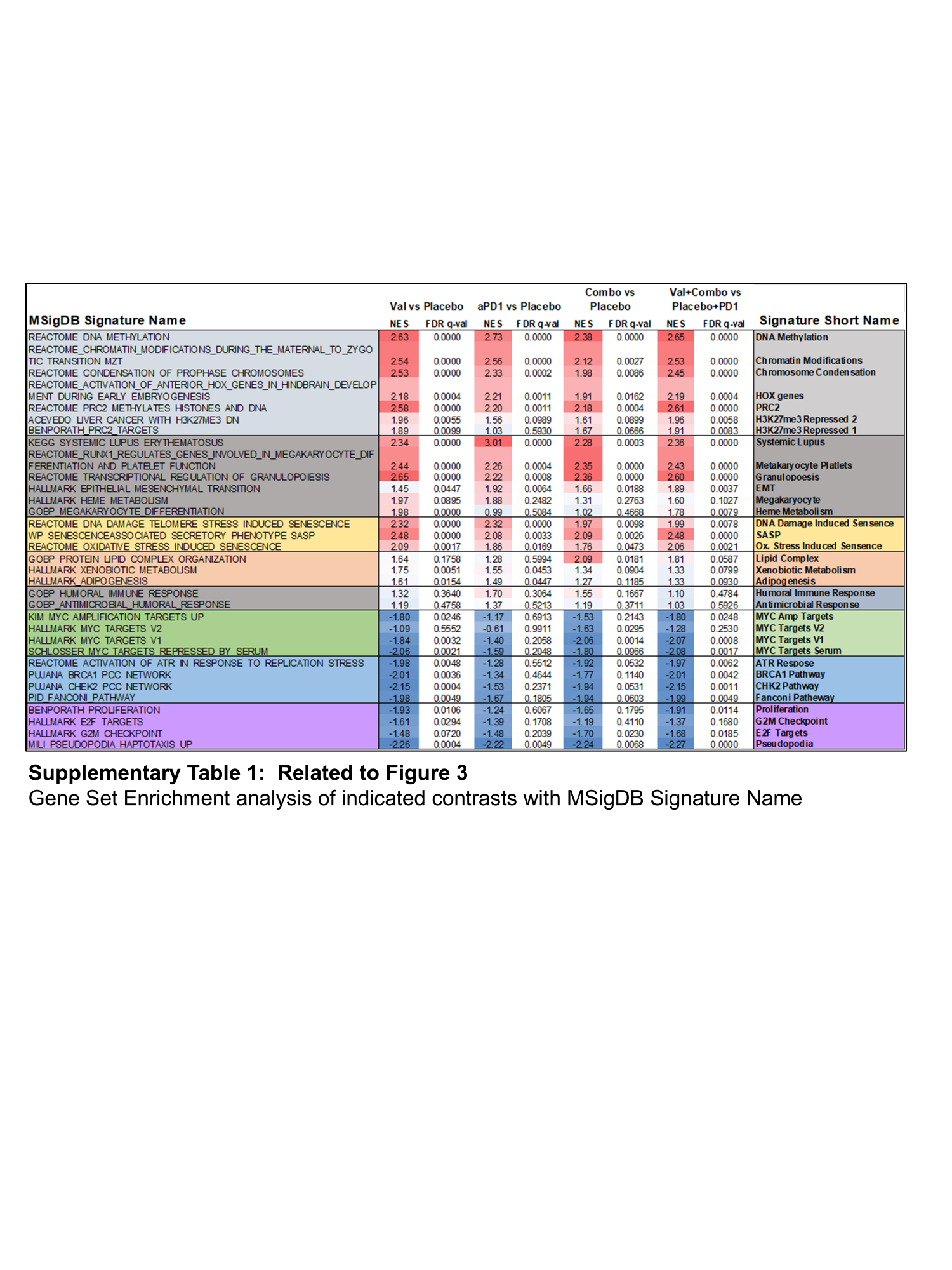

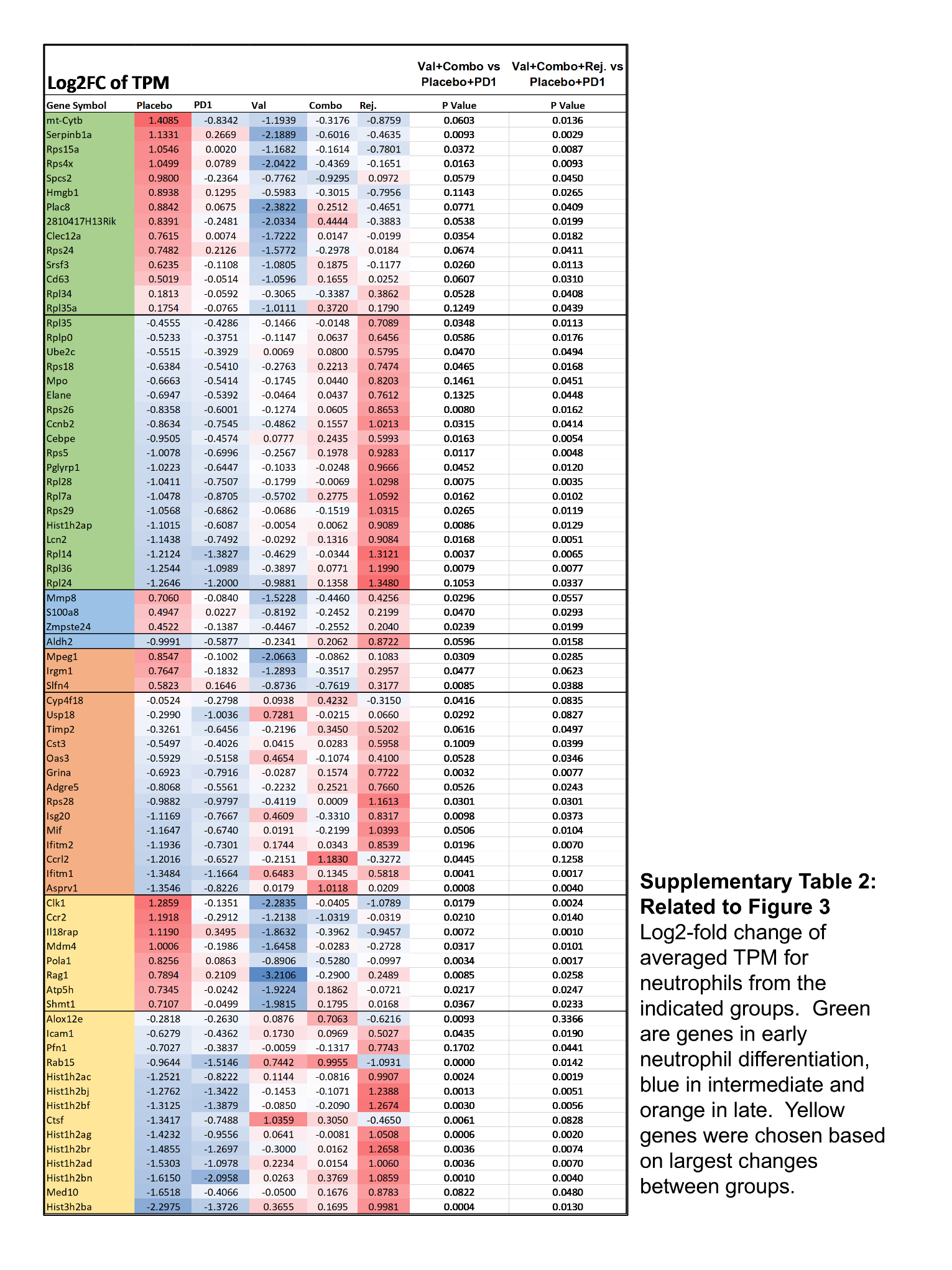

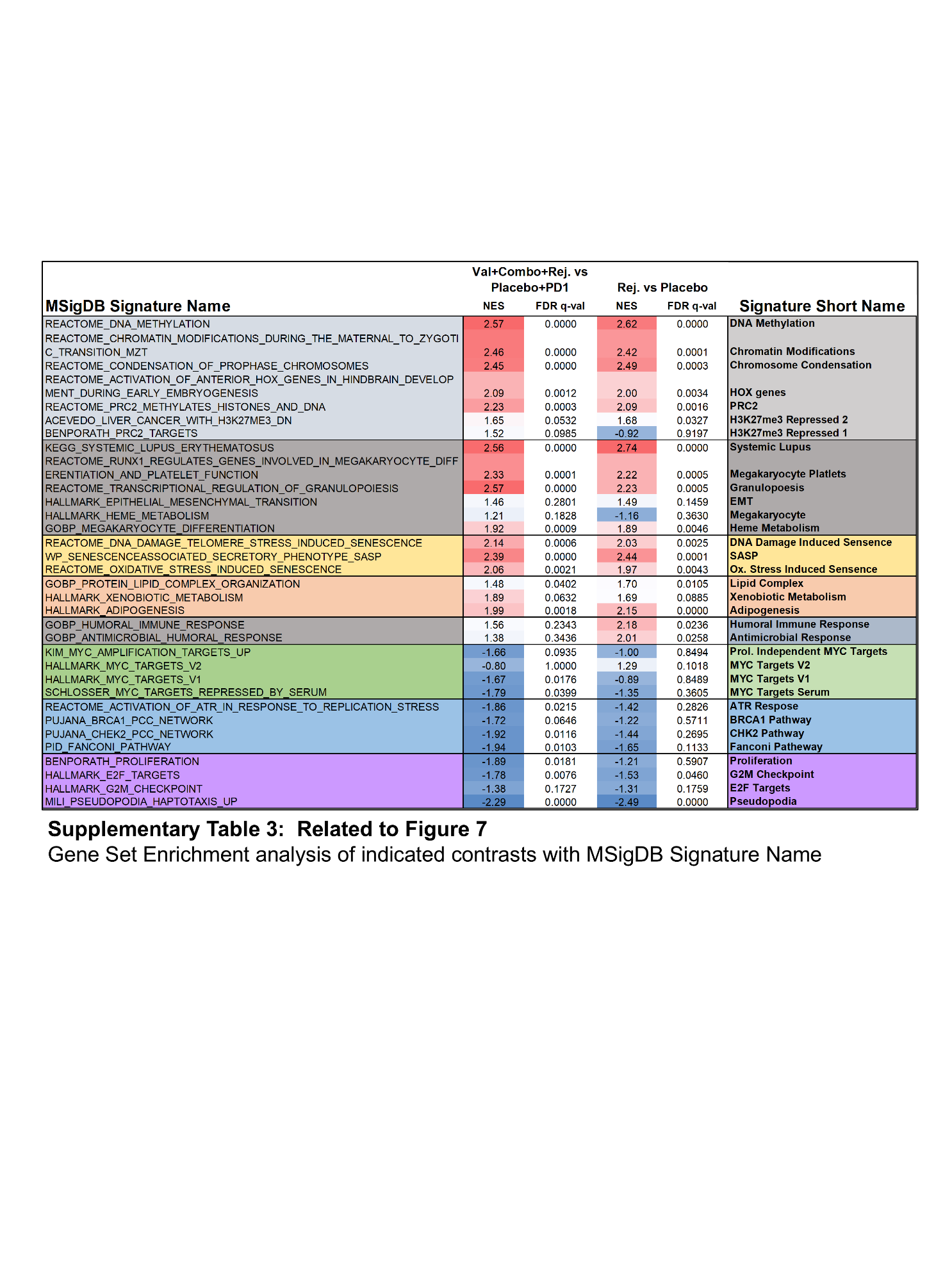
